# Urbanization as an environmental filter for megacolorful birds

**DOI:** 10.1101/2024.06.24.600482

**Authors:** Lucas Ferreira do Nascimento, Paulo R. Guimarães, Julian Evans, W. Daniel Kissling

## Abstract

The colorfulness of bird plumage plays a crucial role in intraspecific (e.g. sexual display) and interspecific ecological interactions (e.g. camouflage and predation). Consequently, bird plumage colorfulness can affect the success of individuals in novel environments, such as urban settings. However, our understanding of the impact of urbanization on the plumage colorfulness of birds, especially in tropical regions, is limited. To address this gap, we analyzed whether urban environments serve as environmental filters for plumage colorfulness in passerine (Passeriformes) bird assemblages across the biomes of Brazil, the world’s largest tropical country. Using generalized linear models that incorporate bird checklists, functional traits, and a continuous urbanization metric, we show that urbanization increases specific traits that are associated with plumage colorfulness in bird assemblages (i.e. proportion of omnivores, proportion of larger species, and average sexual dichromatism). While the average colorfulness of bird assemblages did not change with increasing urbanization, a negative correlation between the presence of megacolorful birds (i.e. the 5% most colorful species) and urbanization was detected, particularly in biomes with high urban concentrations, such as the Atlantic Forest and the Caatinga. This suggests that urban environments can be unsuitable for the most colorful tropical bird species. Our study additionally shows that factors like body size, diet, and sexual dichromatism play a mediating role in the urban filtering process. Our analyses provide insights into how urban environments act as environmental filters and can help to better understand the consequences of urbanization for tropical biodiversity.

## 1. Introduction

Urbanization is a major threat to biodiversity worldwide (Grimm et al. 2008). Estimates based on night-time lights and artificial impervious areas show that urban land cover has increased over the last 30 years and now covers 0.5–0.7% (ca. 800,000 km^2^) of the total land surface on Earth (Gong et al. 2020, Zhao et al. 2022). Moreover, global projections of urban land cover over the 21st century suggest that the total amount of urban land will further increase by a factor of 1.8–5.9, with the fastest urban land expansion occurring in the tropics (Seto et al. 2012, Gao et al. 2020). Urbanization has a strong impact on biodiversity through the loss of natural habitats and through filtering certain species that cannot survive in urban habitats (Aronson et al. 2016, Hoang & Kanemoto 2021). Urbanization can thus alter the composition of species assemblages, ecological interactions among species, and evolutionary processes such as sexual and natural selection (Hahs & Evans 2015, Iglesias-Carrasco et al. 2019). For instance, recent studies suggest that urban tolerant bird species are more generalist, can have either larger or smaller body sizes, and show less plumage sexual dichromatism compared to bird species that do not tolerate urban environments (Callaghan et al. 2019, Hahs et al. 2023, Iglesias-Carrasco et al. 2019). Understanding why certain species thrive in urban environments while others struggle necessitates an analysis of the functional traits linked to resource utilization and risk mitigation in the face of urbanization (Sol et al. 2014). However, the effects of urbanization on the functional trait composition of species assemblages still remains little explored, particularly in tropical regions and for a wide range of functional traits (Afrifa et al. 2022, Patankar et al. 2021).

One of the functional traits that plays an important role in intraspecific and interspecific communication is animal coloration (Cuthill et al. 2017). Animal coloration depends on the capacity of the observer to recognize the hue, saturation, and intensity of light that is reflected from the surface of an organism. The diversity of intra-individual coloration (‘colourfulness’) allows organisms to be conspicuous and mediate ecological interactions and sexual selection (Renoult et al. 2017). Over the last decades, knowledge about animal color vision and processes underlying the production, selective forces and community patterns of animal coloration has substantially improved (Cuthill et al. 2017). We now know that birds can produce plumage colorations by feather structures and pigments (e.g. melanin and carotenoid) and that bird vision can be reliably assessed with a tetrahedral color space model (Bostwick 2016, Stoddard & Prum 2008). Recent studies have found that plumage colourfulness of passerine bird species (Passeriformes) is mainly driven by sexual dichromatism, by diets that are rich in carotenoid pigments (fruits and nectar), and by body size and environmental factors such as temperature, precipitation and forest availability (Cooney et al. 2022). These findings shed light on how various factors plumage colourfulness in bird assemblages, but the role of urbanization as an environmental filter for the plumage colourfulness of bird assemblages remains little studied (Patankar et al. 2021). Understanding how bird plumage colorfulness mediates the success of birds in urban environments can assist in planning more biodiverse cities amid the expansion of urban areas.

In urban environments, dietary constraints and predation risk could be two key factors for filtering the colourfulness in bird assemblages (Leveau 2019, Yu et al. 2024). Dietary constraints might play a role if urban environments favor omnivorous species and reduce birds that have very specialized feeding strategies, such as frugivores, nectarivores, insectivores and granivores (Callaghan et al. 2019, Gutiérrez-Tapia et al. 2018). Omnivorous birds can make use of food resources that are discarded by humans (Patankar et al. 2021) and birds with specialized feeding strategies might have restricted food resources in urban environments (Gutiérrez-Tapia et al. 2018). Omnivorous species are thus more likely to survive in environments with high levels of urbanization than specialized ones. Moreover, changes in background colors from green in natural areas with low urbanization to gray in heavily urbanized areas (Hilal 2018) can increase the predation risk of conspicuous animals in urban environments (Delhey & Peters 2017, Leveau 2019). Predation risk may also be related to body size, with small passerine birds having a higher predation risk than large-bodied species (Götmark & Post 1996). Within passerine birds, small species tend to be more colorful than large ones (Cooney et al. 2022). This might increase the predation risk of small birds in urban environments, so that large passerine birds in cities might have a larger chance of surviving. We thus expect a negative association between urbanization and plumage colourfulness of passerine bird assemblages if limited food supply and high predation risk reduces the chance of colorful birds to survive in urban environments.

Pressures on sexual selection might also affect the success of colorful bird species in urban environments. Organisms in urban environments are exposed to novel abiotic pressures such as changes in climatic conditions, soil chemistries, hydrological processes, and nutrient dynamics, or acoustic and light pollution (Grimm et al. 2008). These pressures might change the costs and benefits arising from sexual selection and influence the establishment of species in urban environments, as organisms must invest in sexual ornaments (e.g., singing and coloration) while dealing with the challenges posed by environmental changes (Candolin & Heuschele 2008). For example, the males of a Neotropical frog species have increased the conspicuousness of their calls from forest to urban environments, but this has not led to an increase in their sexual attractiveness to females (Halfwerk et al. 2019). In contrast, the decline in carotenoid-based coloration of male house finches in North America along a gradient of urbanization is associated with a change in female color preferences, suggesting an influence of urbanization on female mate-choice behavior (Giraudeau et al. 2018). A widely used measure of sexual selection in birds is plumage sexual dichromatism which reflects the plumage color differences between males and females (Dale et al. 2015). Phylogenetic comparative analyses have shown that plumage sexual dichromatism of passerine birds can be negatively associated with their tolerance to live in urban environments (Iglesias-Carrasco et al. 2019). We can thus expect a negative association between urbanization and plumage colourfulness of passerine bird assemblages if sexual selection reduces the chance of colorful birds to survive in urban environments.

Here, we test to what extent urban environments can act as an environmental filter on plumage colourfulness of passerine assemblages. We conduct our study across Brazil, the largest tropical country in the world, using bird checklists from a large-scale citizen science project (Sullivan et al. 2009), comprehensive functional trait information from novel databases (Cooney et al. 2022, Tobias et al. 2022) and harmonized land cover information from different biomes (Souza et al. 2020). Since various functional traits might mediate the relationship between plumage colourfulness and urbanization, we first test whether relationships exist between urbanization and omnivorous diets, body size, and sexual dichromatism. In a second step, we quantify the effect of urbanization on the average colourfulness of bird assemblages and on the presence of species with the highest plumage colourfulness values (‘megacolorful birds’) (see examples in Figure 1). We control for confounding factors and test for interactions with diet, body size, and sexual dichromatism, and whether those relationships vary by biome. We expect a positive relationship between urbanization and the proportion of omnivores and larger bird species within species assemblages (Figure 2a–b) and a negative relationship between urbanization and average sexual dichromatism (Figure 2c). Moreover, we expect a negative relationship between urbanization and passerine plumage colourfulness (Figure 2d), especially for the megacolorful birds. Our analyses provide insights into the role of urban environments as environmental filters and can help to better predict the consequences of urbanization for tropical biodiversity (Aronson et al. 2016, Hahs & Evans 2015).

**Figure 1.**
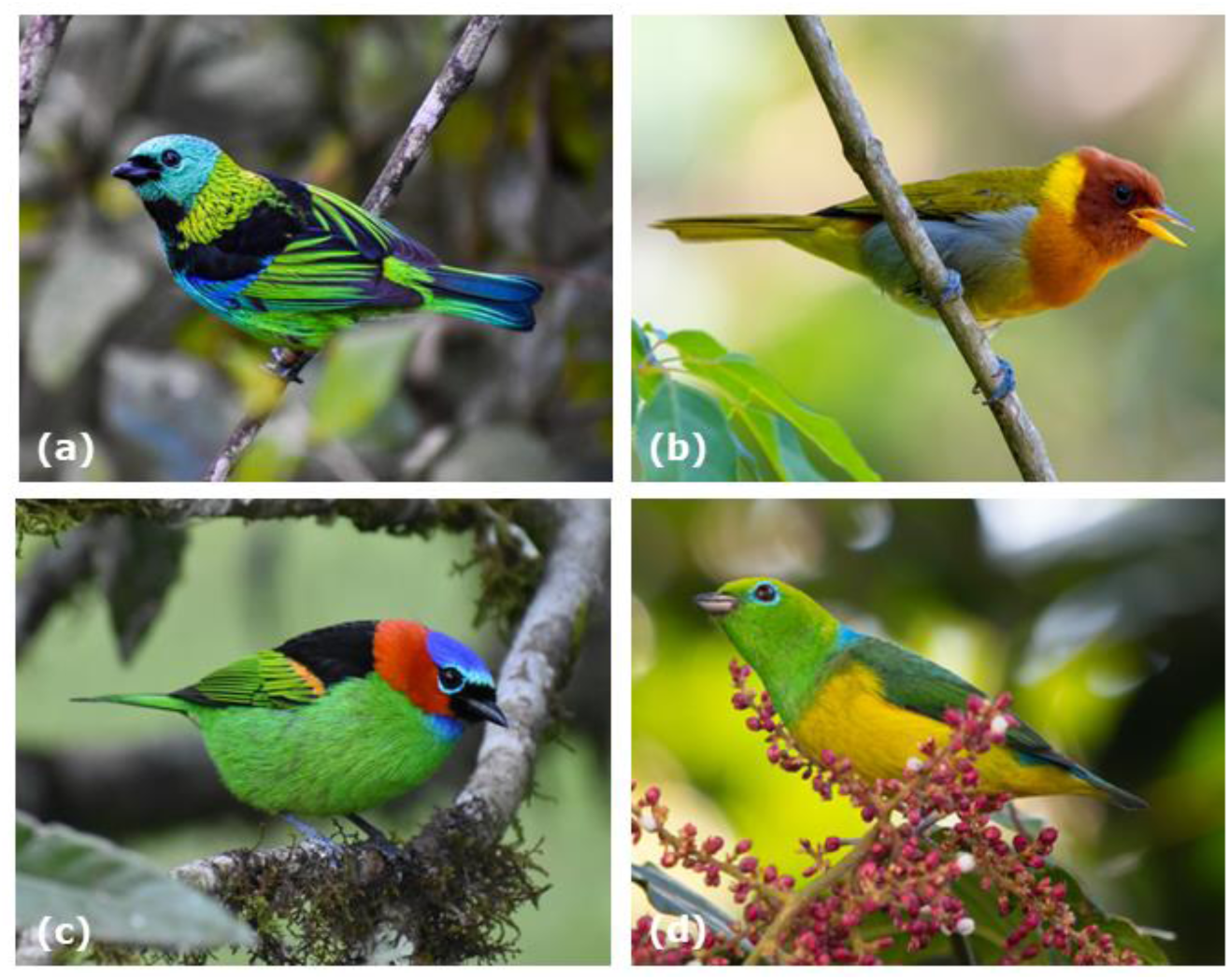
Examples of megacolorful bird species. (a) Green-headed Tanager (*Tangara seledon*, Thraupidae), (b) Rufous-headed Tanager (*Hemithraupis ruficapilla*, Thraupidae), (c) Red-necked Tanager (*Tangara cyanocephala*, Thraupidae), and (d) Blue-naped Chlorophonia (*Chlorophonia cyanea*, Fringillidae). Pictures by Mathias M. Pires.

**Figure 2.**
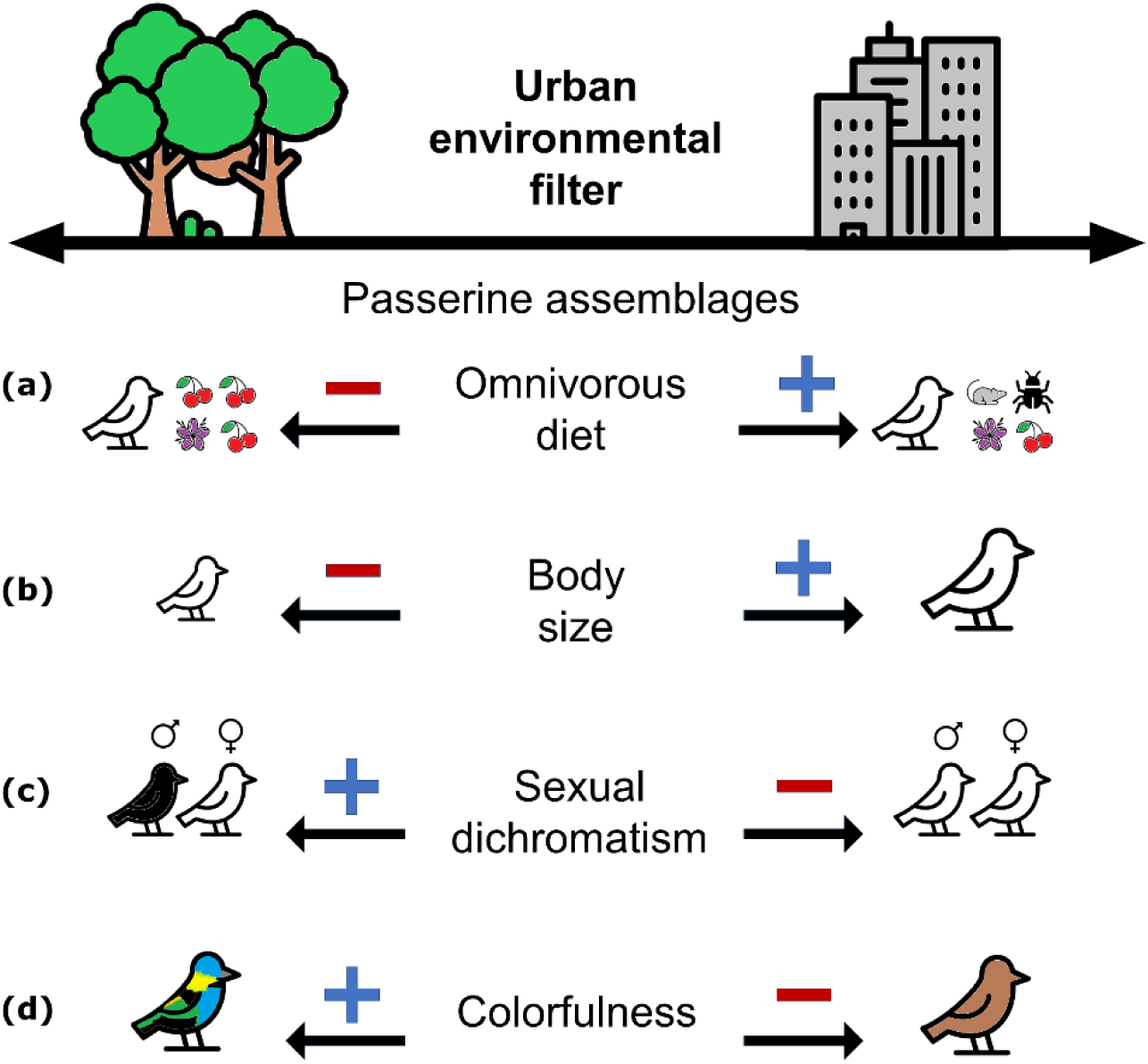
Urbanization as an environmental filter on the representation of functional traits (diet, body size, sexual dichromatism) and colorfulness of passerine birds. Urbanization is expected to increase the proportion of (a) omnivorous passerines and (b) large-bodied species, and decrease (c) average sexual dichromatism and (d) colorfulness of passerine assemblages.

## 2. Material and methods

We designed a workflow to test whether urban environments act as an environmental filter on the plumage colorfulness of passerine assemblages across Brazil (Figure 3). To do so, we integrated different sources of raw data (Figure 3a–c) to quantify functional and environmental characteristics of passerine assemblages (Figure 3d) and used these variables to statistically analyze the effect of urbanization on functional traits and colorfulness of birds in different biomes (Figure 3e).

**Figure 3.**
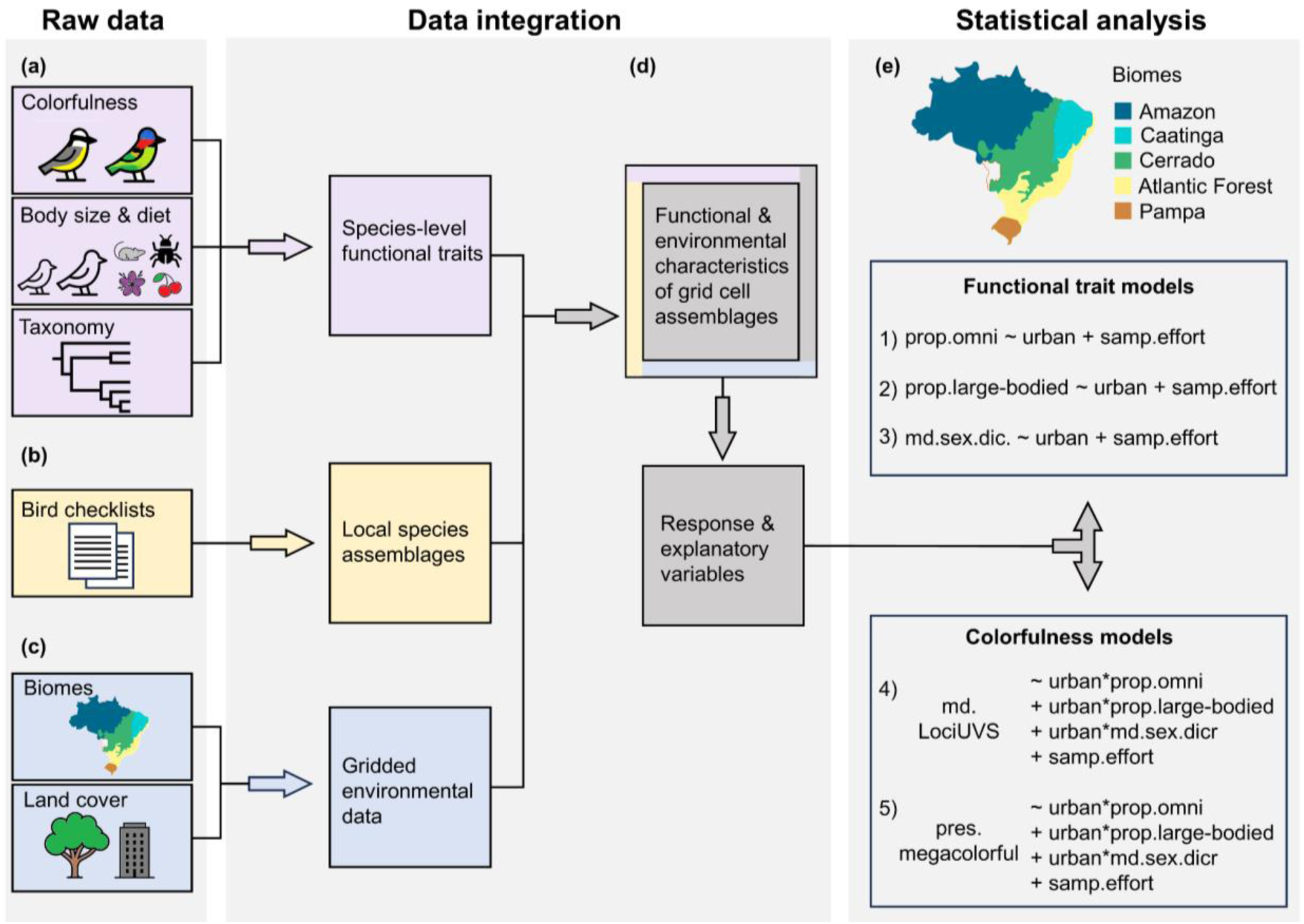
Workflow for integrating various raw datasets to perform statistical analyses of urbanization on functional traits and colourfulness of bird assemblages. (a) Raw data on colourfulness, body size, diet and taxonomy are integrated into a species-level functional trait dataset. Combined with (b) checklists of bird species assemblages and (c) gridded environmental data, the (d) response and explanatory variables of grid cell assemblages can be generated. The integrated data are used in (e) statistical models to explain functional traits and colourfulness with urbanization and other assemblage characteristics for each biome, taking sampling effort (number of checklists) into account. Abbreviations: urban = urbanization (proportion of urban cells in a radius of 250 m), prop. = proportion, omni = omnivorous species, sex.dic. = sexual dichromatism, pres. = presence, md. = median, samp.effort = sampling effort, LociUVS = number of color loci in an avian UV-sensitive (UVS) tetrahedral color space.

### 2.1 Raw data

We first used a dataset for which the colorfulness of each bird species was quantified using digital images of bird specimens from museum collections (Cooney *et al*. 2022). We used LociUVS as the metric, i.e. the number of color loci in an avian UV-sensitive (UVS) tetrahedral color space (Cooney *et al*. 2022). To calculate the LociUVS metric, RGB values of bird images are converted in a bird vision tetrahedral color space model, where each vertex represents one of the four types of bird eye cones sensitive to long, medium, short, and ultraviolet wavelengths (Stoddard & Prum 2008). The tetrahedral color space is then subdivided in three-dimensional cells termed ‘color loci’. The LociUVS (i.e. colorfulness) of a particular bird is defined by the total number of color loci occupied in the tetrahedral color space, i.e. the diversity of bird coloration taking bird vision into account. We extracted the LociUVS data of all passerine bird species from Cooney *et al*. (2022). We primarily used the male LociUVS data, but also used the female LociUVS data to calculate the sexual dichromatism for each species (i.e. the absolute difference between the LociUVS of males and the LociUVS of females).

To quantify the diet and body size of passerine bird species, we used the diet categorization and body mass data from AVONET (Tobias et al. 2022). AVONET captures information from literature, fieldwork and museum specimens and is the most comprehensive, up-to-date functional trait dataset of birds worldwide. We categorized the diet of each species as either omnivorous (dietary generalist) or not (dietary specialist). The AVONET diet dataset provides the percentage of food items in different diet categories and defines omnivores as species utilizing multiple food items in relatively equal proportions (all lower than 60%). In contrast, specialized species are species that predominantly feed on one diet category, i.e. frugivores (feeding mainly on fruit), granivores (feeding mainly on seeds or nuts), nectarivores (feeding mainly on nectar), herbivores (feeding mainly plant materials such as leaves and buds), invertivores (feeding mainly on invertebrates), and aquatic predators (feeding mainly on aquatic vertebrates and invertebrates). For body size, data of AVONET (average body mass in g per species) were used to classify species into “large” or “small” relative to all species in each biome. This was done by analyzing the frequency distribution of body masses for all species in a biome and classifying species in the upper quartile as “large” and those in the lower quartile as “small” (Figure S1).

We used bird checklists from eBird (Sullivan et al. 2009) to characterize local passerine species assemblages across Brazil. The data are collected with a large-scale citizen science project (Sullivan et al. 2009) and represent a list of bird species seen or heard in a given location. For each checklist, volunteer birdwatchers additionally provide survey information such as date, time, latitude, longitude, whether checklists are complete, and survey type (stationary, traveling or casual observation). We only used stationary surveys to avoid casual observations and transect routes going through different land cover types. To further minimize bias, we followed the eBird best practices guide (Strimas-Mackey et al. 2020) and only included complete checklists that have been reviewed and accepted by eBird specialist reviewers. The eBird basic dataset for Brazil (available at https://ebird.org/home) was downloaded in November 2022. We exclusively used the checklists from the year 2021 as this provided the largest number of checklists in a given year (Figure S2) and the most recent information on bird distributions across the sites.

To quantify urbanization and biome distributions in Brazil, we used the land cover information dataset provided by the MapBiomas project (brasil.mapbiomas.org/en/project). This project provides a cell-per-cell land cover classification at 30 × 30 m resolution, produced from Landsat satellite images and machine learning algorithms through the Google Earth Engine platform (Souza et al. 2020). Each cell has a classification based on the predominance of five major classes (forest, non-forest natural formation, farming, non-vegetated areas, and water) and subclassification levels such as agriculture and urban area. To classify a cell as urban, the algorithm takes non-vegetated surfaces, roads, highways, constructions, human population size and night-time lights into account. We utilized the MapBiomas land cover classification of Brazil from the year 2021 because this matched the year of the bird checklists. The dataset included a biome classification as defined by experts of IBGE - the Brazilian Institute of Geography and Statistics (https://www.ibge.gov.br/apps/biomas/#/home) for the six biomes of Brazil, namely Pantanal, Atlantic Rainforest, Cerrado, Pampa, Amazon, and Caatinga. The number of eBird checklists per land cover class varied among biomes (Figures S3) and we excluded the Pantanal due to the low number of urban eBird checklists being available (*n* = 4). Land cover data were available from MapBiomas in rasterfiles and the biome ranges were available in polygon shapefiles.

### 2.2 Data integration

Using species names, we first integrated the raw data of colorfulness (LociUVS), diet and body size by generating a species by trait matrix of passerine traits (Figure 3a). The species names in AVONET and eBird datasets follow the Clements taxonomy (Clements et al. 2023), while the colorfulness data follows the BirdTree taxonomy (Jetz et al., 2012). Therefore, before conducting species name matching between the raw data, we updated the species names in the colorfulness dataset according to the Clements taxonomy and checked for synonyms using the Avibase (https://avibase.bsc-eoc.org/) and Birds of the World (https://birdsoftheworld.org) online platforms. For the eBird checklists, we combined multiple checklists within a 30 × 30 m grid cell (land cover resolution) and only retained unique species names. This generated local species assemblages (i.e. communities of bird species observed within grid cells) as a species by grid cell matrix (Figure 3b). To account for variation in sampling effort (i.e. available checklists per grid cell), we used the number of checklists as a covariate (see statistical analyses). For biomes and land cover, we used the gridded environmental data to obtain the biome class and land cover class for each 30 × 30 m grid cell (Figure 3c). To quantify urbanization around grid cells with eBird checklists, we calculated the proportion of urban cells in a radius of 250 m. This radius was chosen as a conservative estimate of the home range size of passerine birds (Haddou et al. 2022). We further calculated for each grid cell the proportion of omnivorous species, the proportion of large-bodied species, the median sexual dichromatism, number of checklists and the presence/absence of megacolorful species. To characterize a megacolorful species, we initially created a frequency distribution of LociUVS values across all species within a biome. Subsequently, we identified the 95th percentile as a threshold. Megacolorful species were defined as those having LociUVS values equal to or higher than the 95th percentile cutoff (Figure S4). Then, we classified each grid cell based on the presence (1) or absence (0) of megacolorful species. All spatial analyses were conducted using the geographic coordinate system World Geodetic System 1984 (EPSG:4326). To prevent the computation of proportions in assemblages with very few species, we excluded grid cells containing less than three species. An overview per biome of the total grid cell assemblages (sample size), minimum and maximum number of species in a cell assemblage, total number of species and families is provided in the supplementary material (Table S1).

### 2.3 Statistical analyses

We performed two groups of models for each biome (Figure 3e, Table 1). In the first group (‘functional trait models’), we used two binomial and one Gaussian generalized linear model (GLM) to test how urbanization is related to the proportion of omnivorous birds, the proportion of large-bodied birds and the median sexual dichromatism, respectively. In the second group (‘colorfulness models’), we used one binomial and one Gaussian GLM to test how colorfulness of species assemblages (i.e. median LociUVS and presence of megacolorful species) is related to urbanization and other potentially confounding factors (proportion of omnivorous species, proportion of large species, median sexual dichromatism). We included interaction terms between urbanization and each of the three confounding factors. To account for sampling effort, we also included the number of checklists as a covariate in the GLMs. To improve normality, we log-transformed the two continuous response variables (i.e. median sexual dichromatism and median LociUVS). We explored multi-collinearity among predictor variables by calculating the Variance Inflation Factor (VIF). No variable showed high collinearity (all VIF < 1). To identify and remove non-essential predictor variables, leading to a more interpretable and potentially more generalizable model, we performed a stepwise model selection based on the Akaike information criterion (AIC). To assess the residual diagnostics of the statistical models, we examined the residuals for over/underdispersion, heteroscedasticity, non-linear relationships, homogeneity of variance, and zero inflation issues. This involved evaluating the relationship between the residuals and theoretical quantiles as well as predicted values. Additionally, we identified outliers and, if necessary, compared results with and without them. Furthermore, to assess the robustness of our findings, we repeated the statistical analysis with two alternative (categorical) classifications of the urbanization variable (replacing the continuous % urbanization). First, we classified each grid cell assemblage into one of four categories based on the proportion of urban cells surrounding the cell assemblage within a 250 m radius: 1) *low*, if it comprised equal to or larger than 0% and smaller than 25%; 2) *intermediate*, if it comprised equal to or larger than 25% and smaller than 50%; 3) *high*, if it comprised equal to or larger than 50% and smaller than 75%; and 4) *very high*, if it comprised equal to or larger than 75%. Second, we implemented a binary classification based on whether the grid cell assemblages had at least one urban cell within a buffer of a 250 m radius or not (1 = yes, 0 = no).

**Table 1.** Response variables, explanatory variables and model structure of the general linear models performed in this study. The *Functional trait models* are models which test how urbanization is related to the proportion of omnivorous birds, the proportion of large-bodied birds and the median sexual dichromatism, respectively. he *Colorfulness models* test how the median LociUVS and presence of megacolorful species is related to urbanization and other confounding factors (proportion of omnivorous species, proportion of large species, median sexual dichromatism). sampling effort was controlled for by using the number of checklists per grid cell. Abbreviations: urban = urbanization (proportion of urban cells in a radius of 250 m), prop. = proportion, omni = omnivorous species, sex.dic. = sexual dichromatism, pres. = presence, md. = median, samp.effort = sampling effort, LociUVS = number of color loci in an avian UV-sensitive tetrahedral color space. Interaction between variables = *.

| Response variable | Explanatory variable |  |  |  |  | Model structure |  |  |
| --- | --- | --- | --- | --- | --- | --- | --- | --- |
|  | urban | urban*<br>omni. | urban*<br>large-bodied | urban*<br>md.sex.<br>dic. | number of<br>checklists | binomial | gaussian | link<br>function |
| Functional trait models |  |  |  |  |  |  |  |  |
| Proportion of omnivorous | X |  |  |  | X | X |  | logit |
| Proportion of large species | X |  |  |  | X | X |  | logit |
| Median sexual dichromatism | X |  |  |  | X |  | X | identity |
| Colorfulness models |  |  |  |  |  |  |  |  |
| Median LociUVS | X | X | X | X | X |  | X | identity |
| Presence of megacolorfuls | X | X | X | X | X | X |  | logit |

All data processing and statistical analyses were conducted using the R programming language version 4.3.0 within the RStudio version 2023.6.0.421 (R Core Team 2023). Data visualization and processing were performed with the “tidyverse” R package (Wickham et al. 2019). Geographical Information System (GIS) operations on raster and shape files were conducted using the packages “terra” (Hijmans 2022), “exactextractr” (Baston 2022) and “sf” (Pebesma & Bivand 2023). Statistical analysis was performed using the packages “usdm” (Naimi et al. 2014) and the R base package “stats”. We explored multi-collinearity among predictor variables using the function *vifcor* of the R package “usdm” and performed the stepwise model selection using the function *step* of the R package “stats”. Moreover, we evaluated the residual diagnostics of the statistical models using the R package “DHARMa” (Hartig 2022).

## 3. Results

### 3.1 Functional trait models

A positive effect of urbanization on the proportion of omnivores, proportion of large-bodied species, and median sexual dichromatism in bird assemblages was revealed, but the effect strength differed among Brazilian biomes (Figure 4). Urbanization was associated with a larger proportion of omnivores in bird assemblages of the Amazon, Atlantic Forest, and Cerrado, but not in the Caatinga and Pampa (Figure 4a, Table S2). Similarly, the proportion of large-bodied species increased with urbanization in bird assemblages of the Amazon, Atlantic Forest, and Cerrado, but not in the Caatinga and Pampa (Figure 4b, Table S3). For the median sexual dichromatism, urbanization also showed a positive relationship in bird assemblages of the Atlantic Forest and Cerrado and additionally in the Pampa, but not in the Amazon and Caatinga (Figure 4c, Table S4). The results remained qualitatively the same when analyzing the relationship between the functional trait compositions and categorical classifications of urbanization for both the four categories and the binary classification (results not shown).

**Figure 4.**
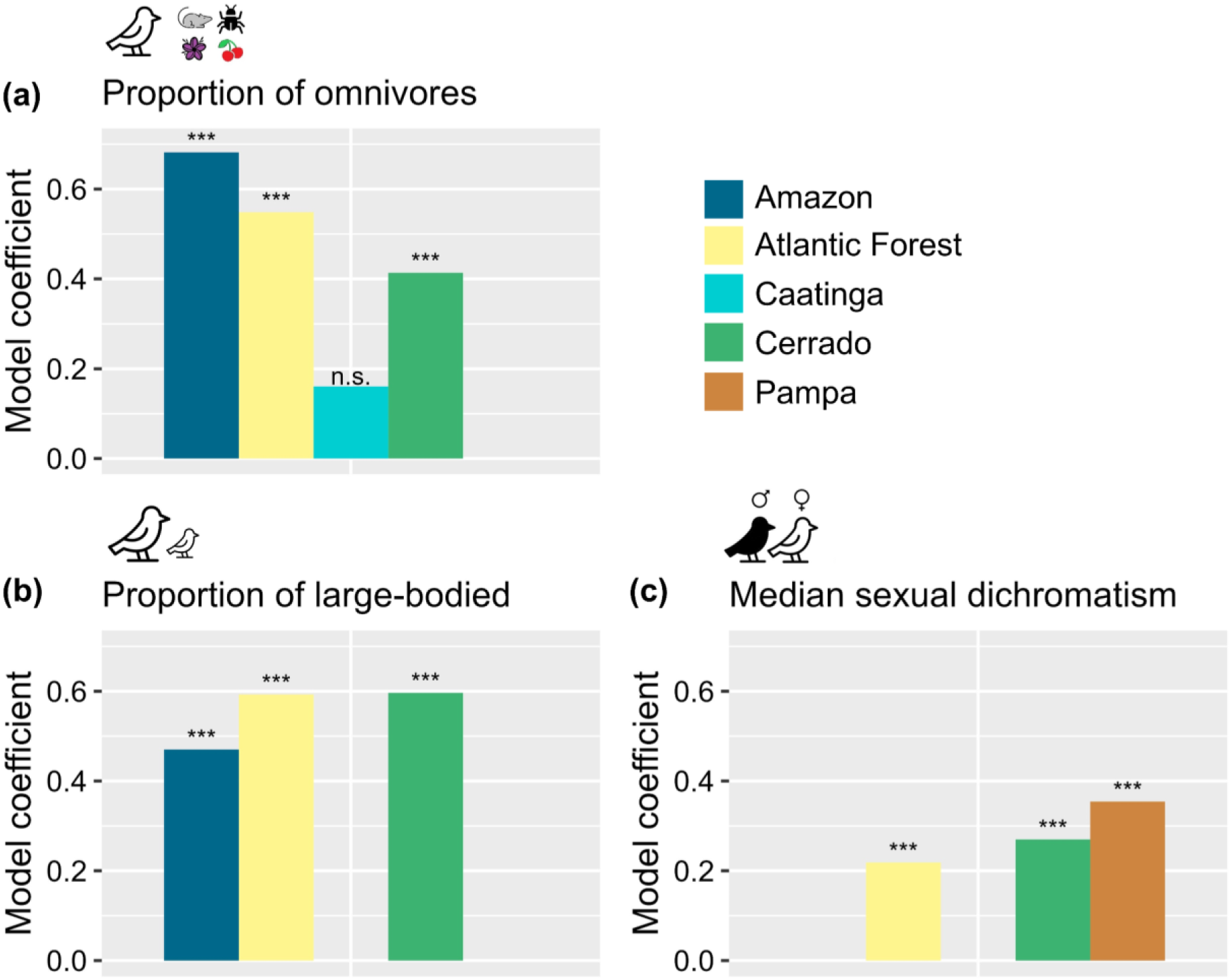
Effects of urbanization on functional trait composition of bird assemblages in Brazilian biomes. Shown are coefficients from the best fitting models for the relationship between urbanization and (a) the proportion of omnivores, (b) the proportion of large-bodied species, and (c) the median sexual dichromatism. See Tables S2–S4 for model details. Significance levels: p < 0.001 = ***, p > 0.05 = n.s.

### 3.2 Colorfulness models

The only statistically significant relationship between urbanization and the median LociUVS was found in the Atlantic Forest, but with a weak effect size (coef. = −0.009, Table S5). Hence, there was no evidence for a strong effect of urban filtering on the average colorfulness of passerine bird assemblages once other traits (proportion of omnivores, proportion of large-bodied species, median sexual dichromatism) and their interactions with urbanization had been accounted for (Table S5). However, there was a relationship between urbanization and the presence of megacolorful species in two biomes of Brazil (Table 2). In the Atlantic Forest and Caatinga, urbanization showed a negative association with the presence of megacolorful species (Figure 5) once other traits and their interactions with urbanization had been accounted for (Table 2). Moreover, the interaction effect between urbanization and the proportion of large-bodied species on the presence of megacolorful birds was statistically significant in the Atlantic Forest, Cerrado, and Caatinga (Table 2), suggesting that assemblages with high proportions of large-bodied species have a stronger decline of megacolorful species along an urbanization gradient than assemblages with small proportions of large-bodied species (Figure 5). Furthermore, the number of checklists per grid cell was positively associated with the presence of megacolorful bird species in bird assemblages across all the Brazilian biomes (Table 2), confirming the importance of controlling for sampling effort. The results remained qualitatively similar when analyzing the relationship between the median LociUVS and presence of megacolorful species in the assemblages using both the four categories and the binary classification of urbanization (results not shown). The taxonomic names of the megacolorful species and the species most frequently recorded in grid cells with high urban coverage are provided in the supplementary material (Table S6 and Table S7).

**Figure 5.**
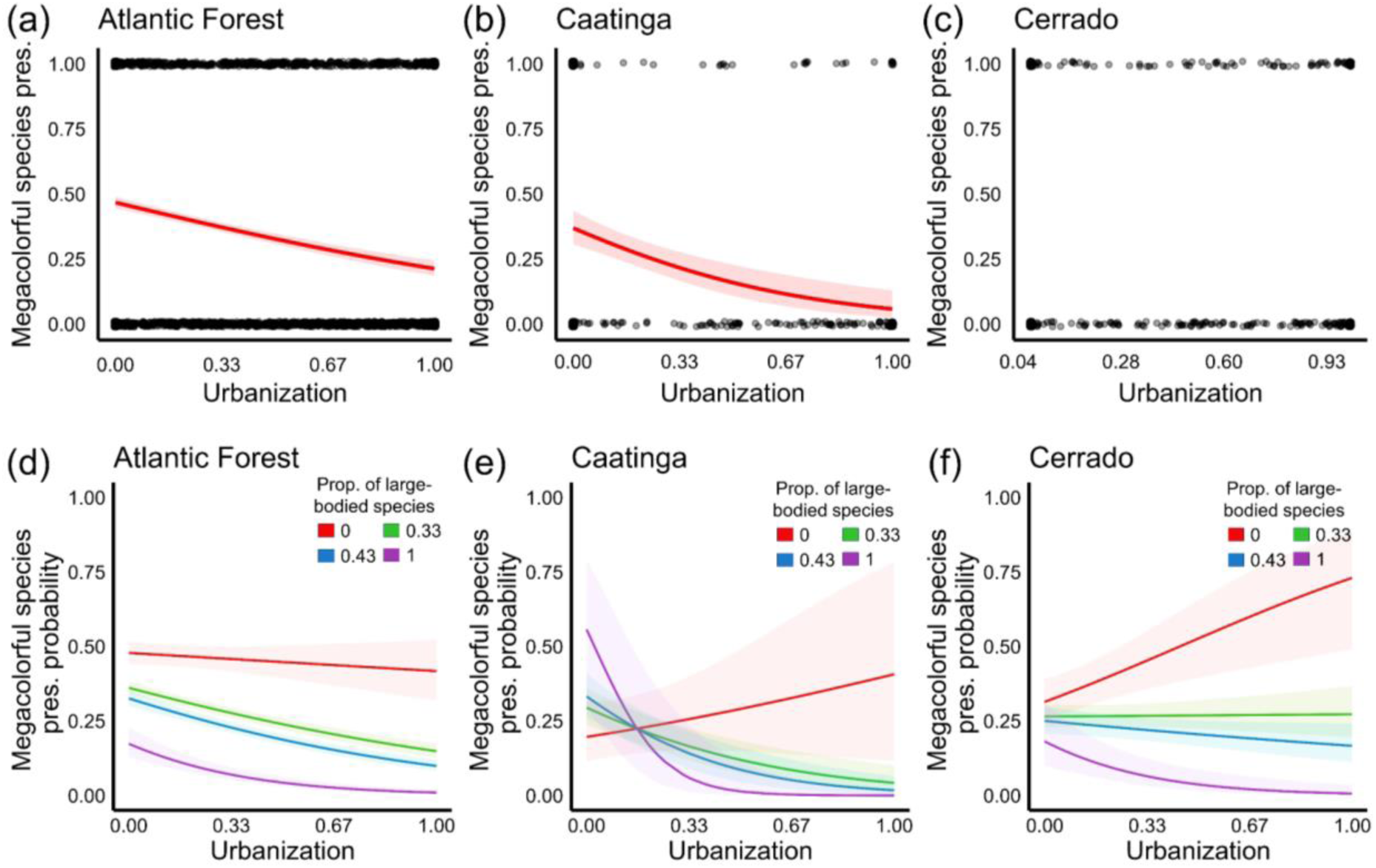
Presence of megacolorful bird species along urbanization gradients in Brazil. Estimated relationships (red lines) between urbanization and the presence-absence of megacolorful species in bird assemblages of (a) the Atlantic Forest, (b) the Caatinga and (c) the Cerrado (no statistically significant relationship). The 95% confidence interval is based on standard errors. When the proportion of large-bodied species in the assemblages increases, the relationship between urbanization and the presence of megacolorful species becomes more negative in (d) the Atlantic Forest, (e) the Caatinga and (f) Cerrado. Abbreviation: pres. = presence. See Table 2 for model details.

**Table 2.**
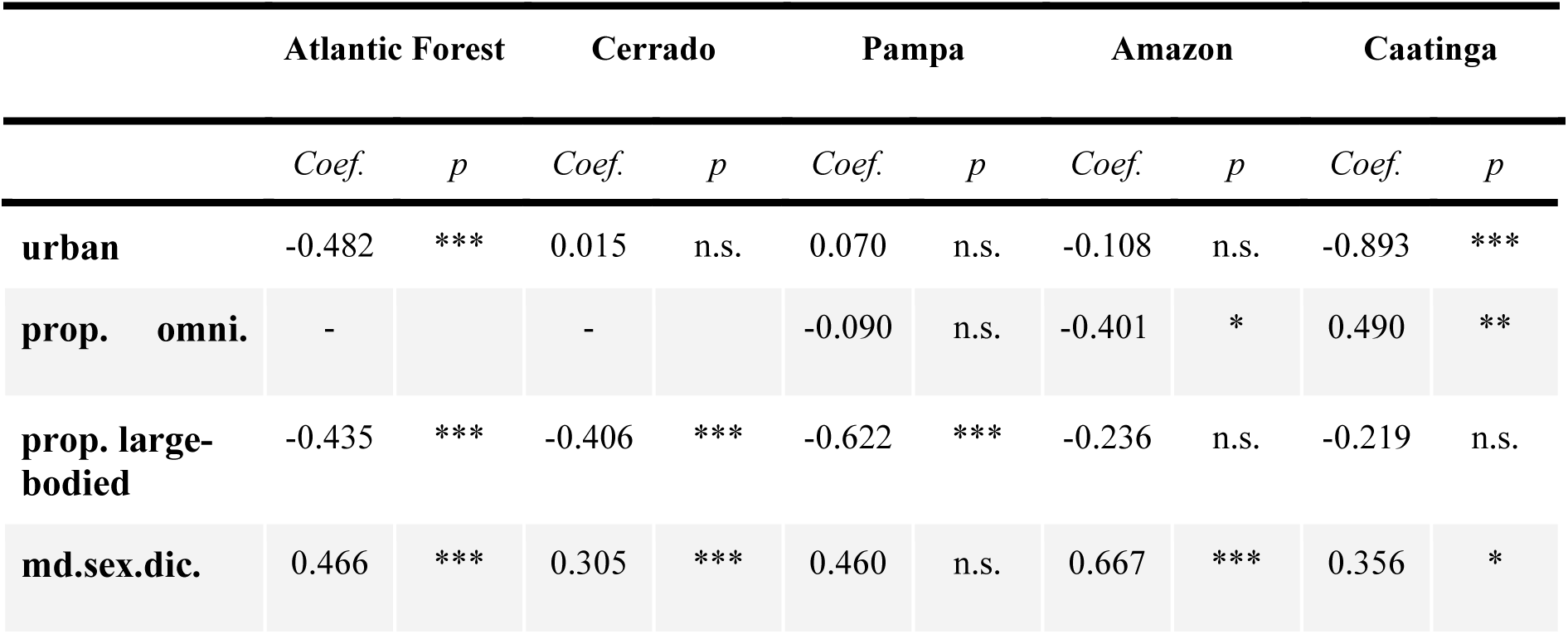
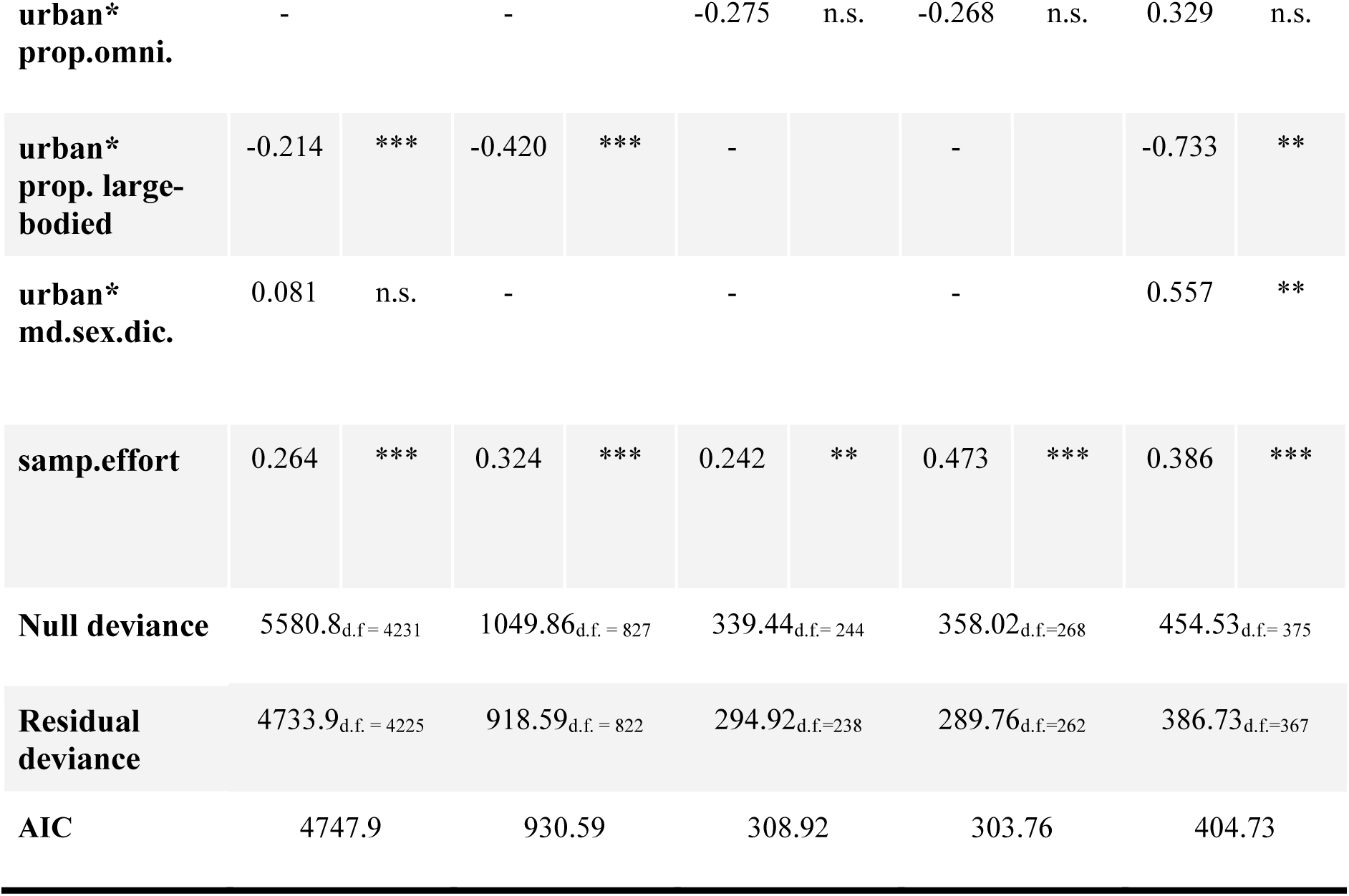
Best fitting model for the relationship between urbanization and the presence of megacolorful species in bird assemblages in different Brazilian biomes. Abbreviations: urban = urbanization, prop. = proportion, omni = omnivorous species, sex.dic. = sexual dichromatism, pres. = presence, md. = median, samp.effort = sampling effort (number of checklists). Interaction between variables = *. Significance levels: p < 0.001 = ***, p < 0.01 = **, p < 0.05 = *, p > 0.05 = n.s.

|  |  | Atlantic Forest |  | Cerrado |  | Pampa |  | Amazon |  | Caatinga |  |
| --- | --- | --- | --- | --- | --- | --- | --- | --- | --- | --- | --- |
|  |  | Coef. | p | Coef. | p | Coef. | p | Coef. | p | Coef. | p |
| **urban** |  | -0.482 | *** | 0.015 | n.s. | 0.070 | n.s. | -0.108 | n.s. | -0.893 | *** |
| **prop.** | **omni.** | - |  | - |  | -0.090 | n.s. | -0.401 | * | 0.490 | ** |
| **prop.** | **large-bodied** | -0.435 | *** | -0.406 | *** | -0.622 | *** | -0.236 | n.s. | -0.219 | n.s. |
| **md.sex.dic.** |  | 0.466 | *** | 0.305 | *** | 0.460 | n.s. | 0.667 | *** | 0.356 | * |

|  |  |  |  |  |  |  |  |  |  |  |
| --- | --- | --- | --- | --- | --- | --- | --- | --- | --- | --- |
| **urban\***<br>**prop.omni.** | - |  | - |  | -0.275 | n.s. | -0.268 | n.s. | 0.329 | n.s. |
| **urban\***<br>**prop. large-**<br>**bodied** | -0.214 | *** | -0.420 | *** | - |  | - |  | -0.733 | ** |
| **urban\***<br>**md.sex.dic.** | 0.081 | n.s. | - |  | - |  | - |  | 0.557 | ** |
| **samp.effort** | 0.264 | *** | 0.324 | *** | 0.242 | ** | 0.473 | *** | 0.386 | *** |
| **Null deviance** | 5580.8<sub>d.f. = 4231</sub> |  | 1049.86<sub>d.f. = 827</sub> |  | 339.44<sub>d.f. = 244</sub> |  | 358.02<sub>d.f. = 268</sub> |  | 454.53<sub>d.f. = 375</sub> |  |
| **Residual deviance** | 4733.9<sub>d.f. = 4225</sub> |  | 918.59<sub>d.f. = 822</sub> |  | 294.92<sub>d.f. = 238</sub> |  | 289.76<sub>d.f. = 262</sub> |  | 386.73<sub>d.f. = 367</sub> |  |
| **AIC** | 4747.9 |  | 930.59 |  | 308.92 |  | 303.76 |  | 404.73 |  |

## 4. Discussion

We found that large-bodied, omnivorous and non-megacolorful passerine species tend to have more success in environments with high urban coverage compared to small-bodied, diet specialists, and megacolorful passerine species across bird assemblages in Brazilian biomes. According to our initial expectations, the proportion of omnivorous and larger passerine species in the assemblages increased with urbanization in Atlantic Forest, Amazon, and Cerrado. Unexpectedly, we also observed a positive relationship between sexual dichromatism and urbanization in the Atlantic Forest, Cerrado, and Pampa. We did not find a significant relationship between urbanization and the average colorfulness of passerine assemblages. However, a decrease in the probability of presence of megacolorful bird species was revealed with increasing urbanization in the Atlantic Forest and Caatinga. Furthermore, in the Atlantic Forest, Cerrado, and Caatinga, an increase in the proportion of larger birds is associated with a decrease in the probability of finding megacolorful birds in highly urbanized areas.

Our results show a strong effect of urban filtering pressures on megacolorful tropical bird species, as well as the role of body size and diet in mediating this urban filtering process. Urban environments may not be suitable habitats for megacolorful passerines because the most colorful birds tend to also have specialized diets and smaller body sizes (Conney et al., 2022). On other hand, our results suggest that urban environments favor omnivorous and large-bodied bird species. Birds with a generalized diet may experience more success than specialized ones because they can forage on more food items including discarded human waste (Hahs et al. 2023, Callaghan et al. 2019). Larger birds in urban environments may experience greater success than smaller ones due to reduced predation risk and increased mobility, allowing them to explore a wider range of resources compared to smaller ones. Moreover, our finding about the interaction effect between urbanization and the proportions of larger birds in the assemblages of the Atlantic Forest, Cerrado and Caatinga underscores the significant role of body size in determining the success of megacolorful passerines in urban environments. Megacolorful birds tend to be small-bodied, but urban environments tend to favor large-bodied, non-megacolorful species.

Other biotic factors, such as the disruption of mutualistic interactions, and abiotic factors, including increased temperature (known as the ‘heat-island effect’) and chemical pollution, could also play a role in determining the presence of colorful birds within urban assemblages. For example, megacolorful birds often engage in foraging within mixed-species flocks, a behavior that enhances foraging efficiency and facilitates the detection of predators. However, this behavior may be compromised in urban environments because colorful passerines often rely on forest habitats with high numbers of co-occurring passerine species (Conney et al., 2022), whereas urban environments typically consist of non-forest habitats with low numbers of co-occurring passerine species (McKinney 2006). Additionally, a recent study suggested that melanic birds (i.e. species with blackish colors) may thrive in urban assemblages due to the protective properties of melanin pigments, which can enhance resistance to tissue abrasion, higher temperatures, and detoxification of toxic metals commonly found in urban air pollution (Turak et al. 2022). Megacolorful birds might thus not thrive in urban environments due to their low resistance to the damaging effects of urban pollution. Future studies could explore how urban abiotic factors impact the success of megacolorful birds in tropical urban environments (Janas et al. 2024). Furthermore, human social factors, such as human hunting activities, could also affect the presence of megacolorful birds in urban environments. Colorful birds are more targeted for wildlife trade (Senior et al. 2022), and the likelihood of bird species being hunted can increase with the proximity to urban areas (Ferreiro-Arias et al. 2024). Future studies could also explore how the interplay between abiotic, biotic and human social factors can affect the presence of megacolorful birds in urban environments.

Our results align with our initial expectations and previous studies which conclude that larger dietary breadth is associated with higher success of birds in urban environments (Hahs et al. 2023, Callaghan et al. 2019). Generalist bird species are more likely than specialists to find suitable environmental, physiological or ecological conditions in urban environments (Bonier et al. 2007). However, this can vary among different terrestrial animal taxa. For instance, in contrast to birds the assemblages of amphibians and reptiles can have a higher proportion of specialists in urban environments (Hahs et al. 2023). On other hand, contrary to our findings, recent global studies found that small-bodied bird species have more success in urban environments than larger ones (Hahs et al. 2023, Neate-Clegg et al. 2023). However, these studies focused on bird species in general, whereas our investigation specifically targeted bird species in the order Passeriformes. Moreover, the difference to our findings could be due to the majority of studies focussing on temperate species assemblages while we investigated exclusively tropical assemblages. Tropical environments feature unique abiotic and biotic conditions which may result in biodiversity patterns and responses differing from those in temperate regions. Hence, our findings suggest that the outcome of the urban filtering process should be studied more widely in tropical environments.

Differences between tropical and temperate assemblages can be also an explanation of our unexpected result of a positive relationship between sexual dichromatism and urbanization in Atlantic Forest, Cerrado and Pampa. These findings suggest that sexual selection does not appear to be negatively affecting the survival of birds in tropical urban assemblages, contrary to findings in temperate urban assemblages (Iglesias-Carrasco et al., 2019). Alternatively, the age of cities can be a determinant of the effects of urbanization filter pressures on biodiversity (Evans et al., 2010). Generally, tropical cities are younger than temperate ones, and consequently, there may not be enough time in tropical cities to observe the negative effects of the interplay between natural selection (caused by urbanization) and sexual selection on tropical passerine assemblages. Moreover, the potential negative effects of urbanization on sexual selection occurring at the population scale may not be strong enough to scale up to the community level which is the scale of our study. Future studies could explore the extent to which sexual dichromatism changes across tropical species populations as they transition from natural to urban environments.

Functional traits can exhibit phylogenetic structure, implying that the urban filtering process may impact species from specific clades more than others (Sol et al. 2017). In total, we identified 45 different megacolorful species, with 27 present in the Atlantic Forest, 16 in the Cerrado, 9 in the Pampa, 28 in the Amazon, and 7 in the Caatinga. These megacolorful species belong to the Cardinalidae, Cotingidae, Fringillidae, Icteridae, Tyrannidae, Pipridae, and Thraupidae families, with the vast majority (71%) belonging to the Thraupidae (24 species) and Pipridae (8 species) families. In contrast, when examining in each biome the fifth most frequently recorded species in checklists within highly urban assemblages, we observed species from 20 different families. The majority (51%) belonged to the Tyrannidae (29 species) and also Thraupidae (29 species) families. These results suggest that certain families may experience the effects of urban filtering pressures more than others, potentially leading to a phylogenetically structured effect of urbanization on tropical megacolorful bird species. However, this should be tested more rigorously in future studies using phylogenetic comparative analyses. For illustration, the Great Kiskadee (*Pitangus sulphuratus*, Tyrannidae), a non-megacolorful, omnivorous, and large-bodied bird species, represents (based on our results) a typical species that can thrive in tropical urban environments. The Great Kiskadee was the most recorded species in highly urbanized environments in the Atlantic Forest and Pampa, the second most recorded in highly urbanized assemblages in the Cerrado and Caatinga, and the third most recorded highly urbanized areas of the Amazon biome.

Variation across biomes may stem from the uneven distribution of urbanization across Brazil, creating potential differences in urban filtering pressures on passerine assemblages. For instance, the Atlantic Forest shows the largest anthropogenic pressures within Brazil (Morellato & Haddad, 2000), concentrating 53% of Brazil’s urban areas in 2021, followed by the Cerrado (23%), Caatinga (12%), Amazon (9%), Pampa (3%), and Pantanal (0.2%) (https://brasil.mapbiomas.org/en/). Hence, variation in urbanization pressures among biomes could lead to differences in the outcomes of urban filtering pressures across biomes. Additionally, urbanization drives biodiversity homogenization by favoring the success of the same urban-tolerant species, resulting in cities from different regions hosting similar species (McKinney 2006). Consequently, as urbanization continues to expand, we can anticipate that bird assemblages in currently less urbanized biomes will eventually resemble those found in highly urbanized biomes in the future. To develop more sustainable and biodiverse cities, it may thus be crucial to understand how functional traits of species help them to adapt to urban environments, especially in hotspots of biodiversity such as the tropics (Patankar et al. 2021, Sol et al. 2014). To our knowledge, this study is the most extensive to date on the impacts of urbanization on the colorfulness of birds in tropical environments.

The unsuitability of urban environments for megacolorful bird species may not only lead to a loss of biodiversity but also a reduction of ecosystem services that these species provide, such as seed dispersal. Seed dispersal is a crucial ecosystem service because it is essential for the reproduction and survival of plants, ensuring the colonization of new habitats and the maintenance of genetic diversity within ecosystems, thus supporting ecosystem health and carbon storage (Bello et al. 2015). Moreover, the loss of megacolorful birds potentially undermines another ecosystem service that is rarely taken into account by studies: the aesthetic value of landscapes (Tribot et al. 2018). The aesthetic value of landscapes plays a crucial role in human well-being, positively influencing individuals’ attitudes towards nature and their engagement in conservation efforts (Tribot et al. 2018). Considering the relationship between the aesthetic value of landscapes and biodiversity is directly relevant to people and could lead efforts in constructing more sustainable cities (Tribot et al. 2018). Furthermore, fostering the connection between urban residents and nature is crucial for conserving natural environments because urbanization often disrupts people’s familiarity and connection with their native biological environment (Dunn et al. 2006, McKinney 2006). Indeed, citizen science has emerged as a powerful tool for people’s interactions with nature, while simultaneously collecting data for addressing ecological questions and monitoring biodiversity (Pocock et al. 2017). Volunteer records have facilitated data-intensive scientific research and monitoring across scales (spatially, temporally, or in terms of data volume) that would otherwise be impractical or too costly to achieve without volunteer participation (Pocock et al. 2017). Promoting citizen science has the potential to enhance our understanding of ecological functioning in biodiversity, enrich people’s aesthetic experiences, and motivate efforts to create biodiversity-friendly cities capable of facing the ongoing biodiversity crisis.

## 5. Conclusions

Our study sheds light on the effect of urban filtering on functional traits of passerine bird assemblages across Brazil. Our findings underscore the strong influence of urban filtering pressures on the most colorful tropical bird species, and highlights the role of body size, diet, and sexual dichromatism in mediating this process. Variation among biomes in urban filtering may stem from differences in the distribution and intensity of urbanization pressures. As urbanization continues to expand, we expect that bird assemblages in currently less urbanized biomes may eventually resemble those found in highly urbanized regions. Urbanization poses a significant threat to biodiversity, often leading to homogenization (e.g. the proliferation of the same urban-tolerant species) and harming ecosystem services (e.g. reducing seed dispersal and the aesthetic value of biodiversity). Promoting citizen science can thus offer a promising avenue for fostering a deeper connection between people and nature while simultaneously advancing our understanding of how urban environments filter biodiversity. To develop more sustainable and biodiverse cities, it will be imperative to understand how functional traits enable species to thrive in urban environments.

## 6. Author contributions

L.F.N. and W.D.K designed the study. L.F.N. and J.E. performed and conducted analyses. L.F.N. and W.D.K. wrote the original draft of the manuscript, and all authors contributed substantially to the final draft.

## 7. Acknowledgements

L.F.N. is funded by a São Paulo Research Foundation PhD scholarship (FAPESP, grants 2020/09286-8 and 2022/04217-3). P.R.G. is funded by Brazil’s Council for Scientific and Technological Development (CNPq; grant 307134/2017-2), and FAPESP (grant 2018/14809-0). We are grateful for the constructive comments and feedback from members of the Biogeography & Macroecology (BIOMAC) lab and the Department Theoretical and Computational Ecology (TCE) at the University of Amsterdam.

## 9. Supplementary Material

**Figure S1.**
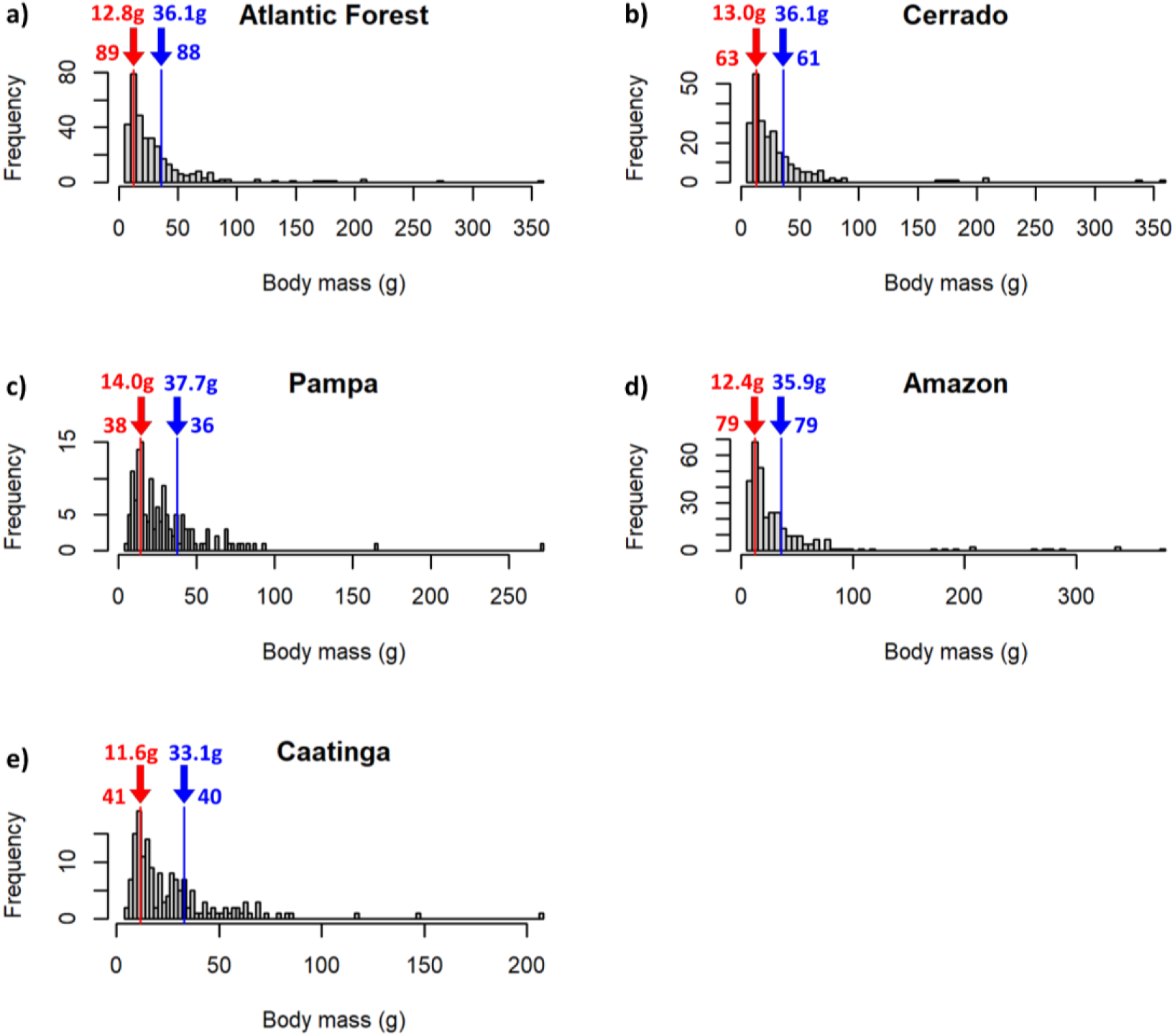
Frequency distribution of species’ body masses (in g) for each biome. Red arrows indicate the first quartile value, while blue arrows indicate the third quartile value. Values above the arrows represent the exact quartile values, while values left and right indicate the number of species with body masses equal to or smaller than the first quartile (red) and equal to or larger than the third quartile (blue).

**Figure S2.**
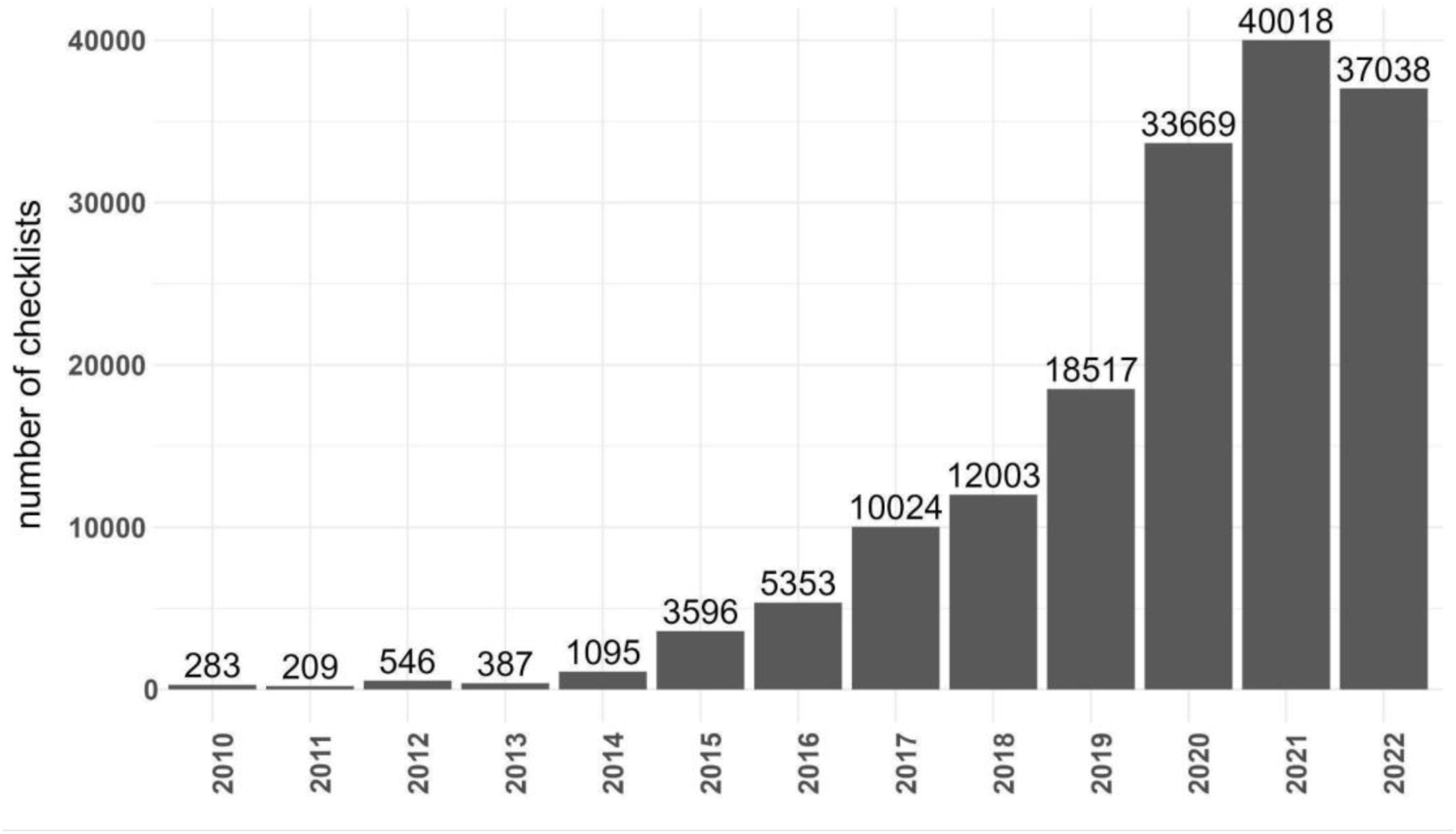
Number of eBird checklists in Brazil by year (downloaded from https://ebird.org/home in November 2022).

**Figure S3.**
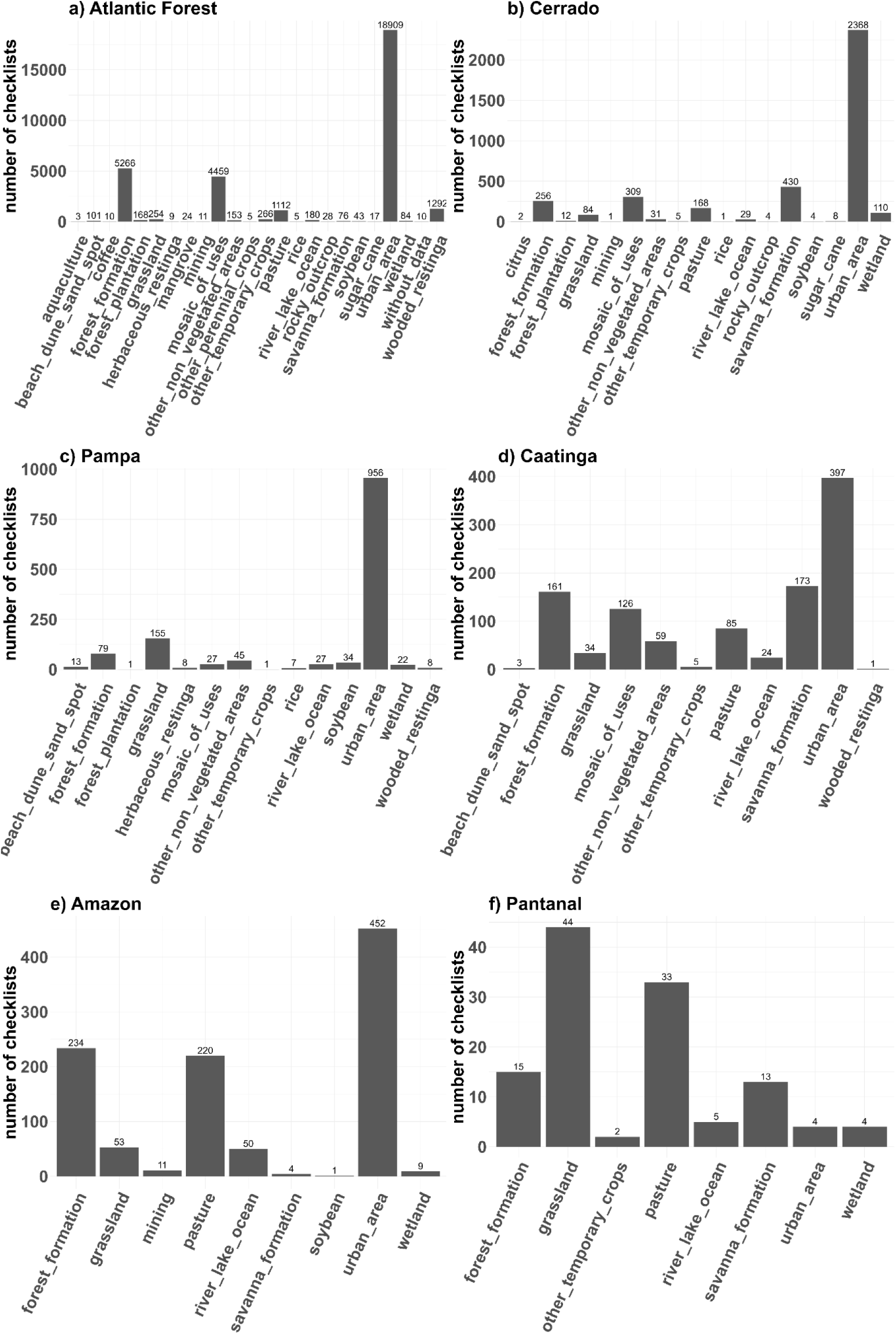
Number of eBird checklists by land use and land cover (MapBiomas) by biome in the year of 2021.

**Figure S4.**
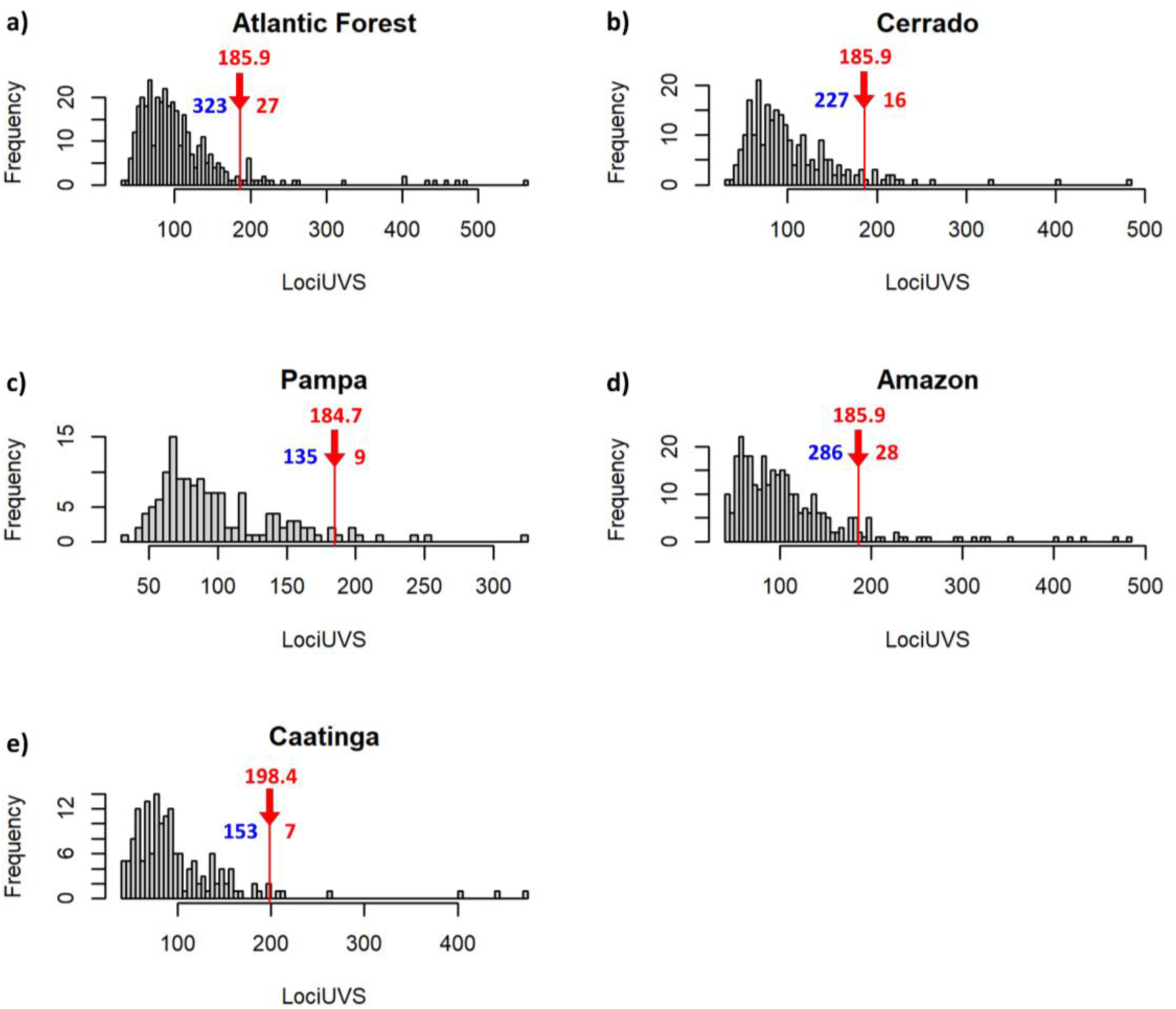
Frequency distribution of species’ LociUVS for each biome. Red arrows indicate 95th percentile value. Values above the arrows (red) represent the exact upper boundary of the 95th percentile value, while values left and right indicate the number of species classified as megacolorful (red) and non-megacolorful (blue).

**Table S1.**
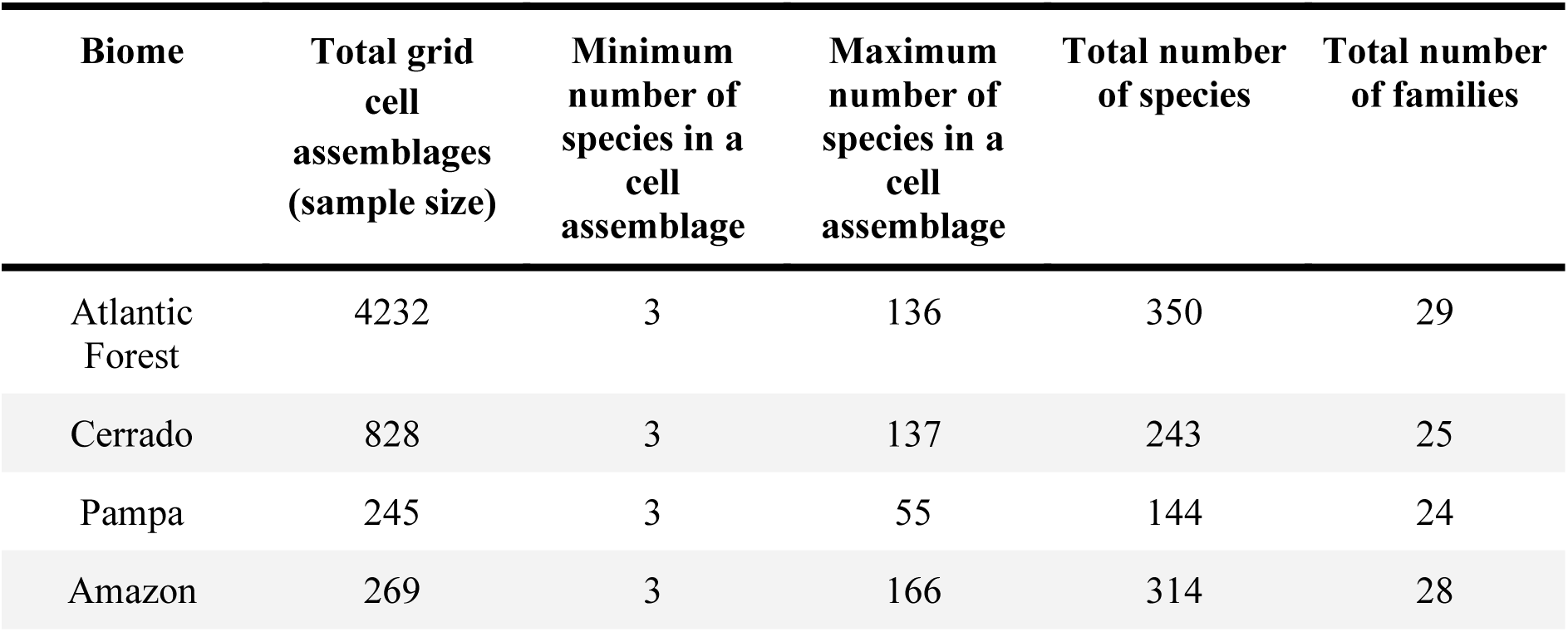
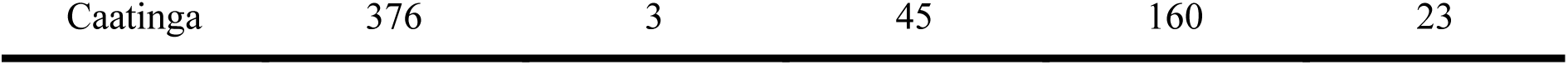
Total grid cell assemblages (sample size), minimum and maximum number of species in a cell assemblage, total number of species and families per biome.

**Table S2.** Proportion of omnivores in relation to urbanization. Coefficient values of the best fitted model estimating the relationship between the proportion of omnivore species in the cell assemblages and urbanization (urban) and sampling effort (number of checklists). Significance levels: p < 0.001 = ***, p < 0.01 = **, p < 0.05 = *, p > 0.05 = n.s.

|  | Atlantic Forest |  | Cerrado |  | Pampa |  | Amazon |  | Caatinga |  |
| --- | --- | --- | --- | --- | --- | --- | --- | --- | --- | --- |
|  | *Coef.* | *p* | *Coef.* | *p* | *Coef.* | *p* | *Coef.* | *p* | *Coef.* | *p* |
| **urban** | 0.548 | *** | 0.413 | *** | - |  | 0.681 | *** | 0.160 | n.s. |
| **samp.effort** | -0.017 | *** | -0.015 | *** | -0.023 | n.s. | -0.030 |  | - |  |
| **Null deviance** | 4394.3 d.f.=4231 |  | 1017.13 d.f.=827 |  | 186.84 d.f.=244 |  | 394.13 d.f.=268 |  | - |  |
| **Residual deviance** | 3917.4 d.f.=4229 |  | 971.62 d.f.=825 |  | 184.02 d.f.=243 |  | 344.33 d.f.=266 |  | 339.89 d.f.=374 |  |
| **AIC** | 12754 |  | 2599.6 |  | 712.17 |  | 930.31 |  | 1066.8 |  |

**Table S3.** Proportion of large-bodied species in relation to urbanization. Best fitting model for the relationship between the proportion of large-bodied species in the cell assemblages and urbanization (urban) and sampling effort (number of checklists). Significance levels: p < 0.001 = ***, p < 0.01 = **, p < 0.05 = *, p > 0.05 = n.s.

|  | Atlantic Forest |  | Cerrado |  | Pampa |  | Amazon |  | Caatinga |  |
| --- | --- | --- | --- | --- | --- | --- | --- | --- | --- | --- |
|  | *Coef.* | *p* | *Coef.* | *p* | *Coef.* | *p* | *Coef.* | *p* | *Coef.* | *p* |
| **urban** | 0.592 | *** | 0.596 | *** | - |  | 0.470 | *** | - |  |
| **samp.effort** | -0.003 | ** | -0.009 | ** | -0.022 | n.s. | -0.017 | *** | - |  |
| **Null deviance** | 5588.2 d.f.=4231 |  | 1282.5 d.f.=827 |  | 198.32 d.f.=244 |  | 292.83 d.f.=268 |  | 414.16 d.f.=375 |  |
| **Residual deviance** | 4968.7 d.f.=4229 |  | 1171.4 d.f.=825 |  | 194.76 d.f.=243 |  | 266.95 d.f.=266 |  | 414.16 d.f.=375 |  |
| **AIC** | 14955 |  | 3068.3 |  | 813.94 |  | 923.12 |  | 1249.4 |  |

**Table S4.**
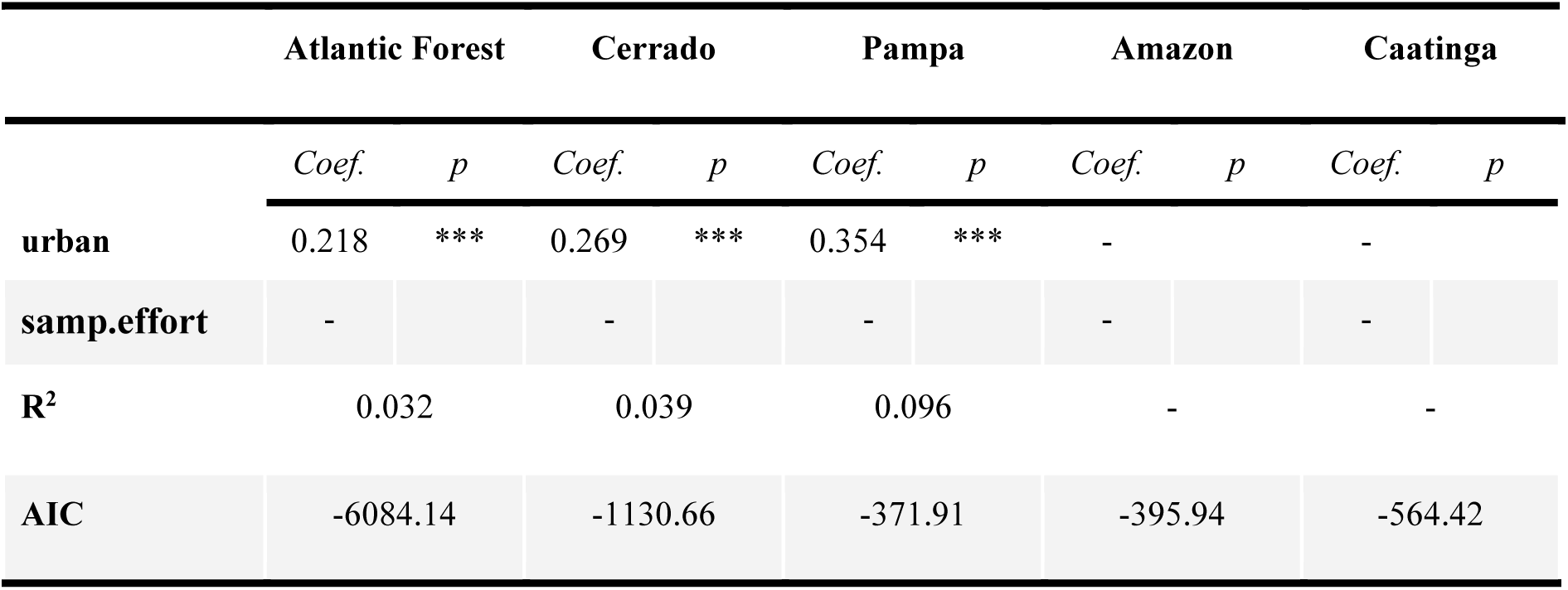
Median of sexual dichromatism in relation to urbanization. Coefficient values of the best fitted model estimating the relationship between the median of sexual dichromatism in the cell assemblages and urbanization (urban) and sampling effort (number of checklists). Significance levels: p < 0.001 = ***, p < 0.01 = **, p < 0.05 = *, p > 0.05 = n.s.

|  | Atlantic Forest |  | Cerrado |  | Pampa |  | Amazon |  | Caatinga |  |
| --- | --- | --- | --- | --- | --- | --- | --- | --- | --- | --- |
|  | *Coef.* | *p* | *Coef.* | *p* | *Coef.* | *p* | *Coef.* | *p* | *Coef.* | *p* |
| **urban** | 0.218 | *** | 0.269 | *** | 0.354 | *** | - |  | - |  |
| **samp.effort** | - |  | - |  | - |  | - |  | - |  |
| **R<sup>2</sup>** | 0.032 |  | 0.039 |  | 0.096 |  | - |  | - |  |
| **AIC** | -6084.14 |  | -1130.66 |  | -371.91 |  | -395.94 |  | -564.42 |  |

**Table S5:** Best fitting model for the relationship between urbanization and the median of LociUVS in grid cell assemblages. Abbreviations: urban = urbanization, prop. = proportion, omni = omnivorous species, sex.dic. = sexual dichromatism, pres. = presence, md. = median, samp.effort = sampling effort (number of checklists). Significance levels: p < 0.001 = ***, p < 0.01 = **, p < 0.05 = *, p > 0.05 = n.s.

|  | Atlantic Forest |  | Cerrado |  | Pampa |  | Amazon |  | Caatinga |  |
| --- | --- | --- | --- | --- | --- | --- | --- | --- | --- | --- |
|  | *Coef.* | *p* | *Coef.* | *p* | *Coef.* | *p* | *Coef.* | *p* | *Coef.* | *p* |
| **urban** | -0.009 | ** | 0.004 | n.s. | -0.008 | n.s. | 0.005 | n.s. | 0.012 | n.s. |
| **prop. omni.** | 0.022 | *** | -0.001 | n.s. | 0.038 | ** | 0.021 | *** | - |  |
| **prop. large-bodied** | -0.029 | *** | -0.008 | n.s. | -0.054 | *** | - |  | - |  |
| **md.sex.dic.** | 0.055 | *** | 0.064 | *** | 0.097 | *** | 0.006 | n.s. | - |  |
| **urban\* prop.omni.** | 0.010 | * | 0.012 | * | -0.016 | n.s. | - |  | - |  |
| **urban\* prop. large-bodied** | 0.039 | * | 0.019 | * | - |  | - |  | - |  |
| **urban\* md.sex.dic.** | - |  | - |  | - |  | 0.040 | * | - |  |
| **samp.effort** | - |  | - |  | - |  | - |  | - |  |
| **R<sup>2</sup>** | 0.117 |  | 0.111 |  | 0.269 |  | 0.109 |  | 0.003 |  |
| **AIC** | -14629.58 |  | -2764.58 |  | -909.6 |  | -864.14 |  | -1378.12 |  |

**Table S6.** Megacolorful species, their respective families, and the biomes where they are present (marked with "X").

| Species name | Family | Atlantic Forest | Cerrado | Pampa | Amazon | Caatinga |
| --- | --- | --- | --- | --- | --- | --- |
| *Granatellus pelzelni* | Cardinalidae |  |  |  | X |  |
| *Cotinga cotinga* | Cotingidae |  |  |  | X |  |
| *Chlorophonia cyanea* | Fringillidae | X |  |  |  |  |
| *Chlorophonia cyanocephala* | Fringillidae | X | X | X |  |  |
| *Euphonia violacea* | Fringillidae | X | X |  | X | X |
| *Euphonia xanthogaster* | Fringillidae | X | X |  | X |  |
| *Euphonia chlorotica* | Fringillidae |  |  | X |  |  |
| *Euphonia finschi* | Fringillidae |  |  |  | X |  |
| *Euphonia rufiventris* | Fringillidae |  |  |  | X |  |
| *Icterus jamacaii* | Icteridae | X | X |  |  | X |
| *Ilicura militaris* | Pipridae | X | X |  |  |  |
| *Machaeropterus regulus* | Pipridae | X |  |  |  |  |
| *Pipra fasciicauda* | Pipridae | X | X |  | X | X |
| *Ceratopipra rubrocapilla* | Pipridae | X |  |  | X |  |
| *Machaeropterus pyrocephalus* | Pipridae |  | X |  | X |  |
| *Lepidothrix iris* | Pipridae |  |  |  | X |  |
| *Ceratopipra erythrocephala* | Pipridae |  |  |  | X |  |
| *Pipra filicauda* | Pipridae |  |  |  | X |  |
| *Chlorophanes spiza* | Thraupidae | X |  |  | X |  |
| *Cyanerpes cyaneus* | Thraupidae | X | X |  | X |  |
| *Dacnis cayana* | Thraupidae | X | X |  | X | X |
| *Hemithraupis ruficapilla* | Thraupidae | X | X |  |  |  |
| *Pipraeidea melanonota* | Thraupidae | X |  | X |  |  |
| *Ramphocelus bresilius* | Thraupidae | X |  | X |  |  |
| *Sicalis flaveola* | Thraupidae | X | X | X | X |  |
| *Stilpnia cayana* | Thraupidae | X | X |  | X | X |
| *Tangara cyanocephala* | Thraupidae | X |  |  |  | X |
| *Tangara cyanoventris* | Thraupidae | X |  |  |  | X |
| *Tangara desmaresti* | Thraupidae | X |  |  |  |  |
| *Tangara mexicana* | Thraupidae | X | X |  | X |  |
| *Stilpnia preciosa* | Thraupidae | X |  | X |  |  |
| *Tangara seledon* | Thraupidae | X |  |  |  |  |
| *Tangara velia* | Thraupidae | X |  |  | X |  |
| *Rauenia bonariensis* | Thraupidae | X | X | X |  |  |
| *Stilpnia cyanicollis* | Thraupidae |  | X |  | X |  |
| *Cyanerpes caeruleus* | Thraupidae |  |  |  | X |  |
| *Cyanerpes nitidus* | Thraupidae |  |  |  | X |  |
| *Tangara chilensis* | Thraupidae |  |  |  | X |  |
| *Tangara gyrola* | Thraupidae |  |  |  | X |  |
| *Stilpnia nigrocincta* | Thraupidae |  |  |  | X |  |
| *Ixothraupis punctata* | Thraupidae |  |  |  | X |  |
| *Ixothraupis varia* | Thraupidae |  |  |  | X |  |
| *Piprites pileata* | Tyrannidae | X |  |  |  |  |
| *Pyrocephalus rubinus* | Tyrannidae | X | X | X | X |  |
| *Tachuris rubrigastra* | Tyrannidae |  |  | X |  |  |

**Table S7.**
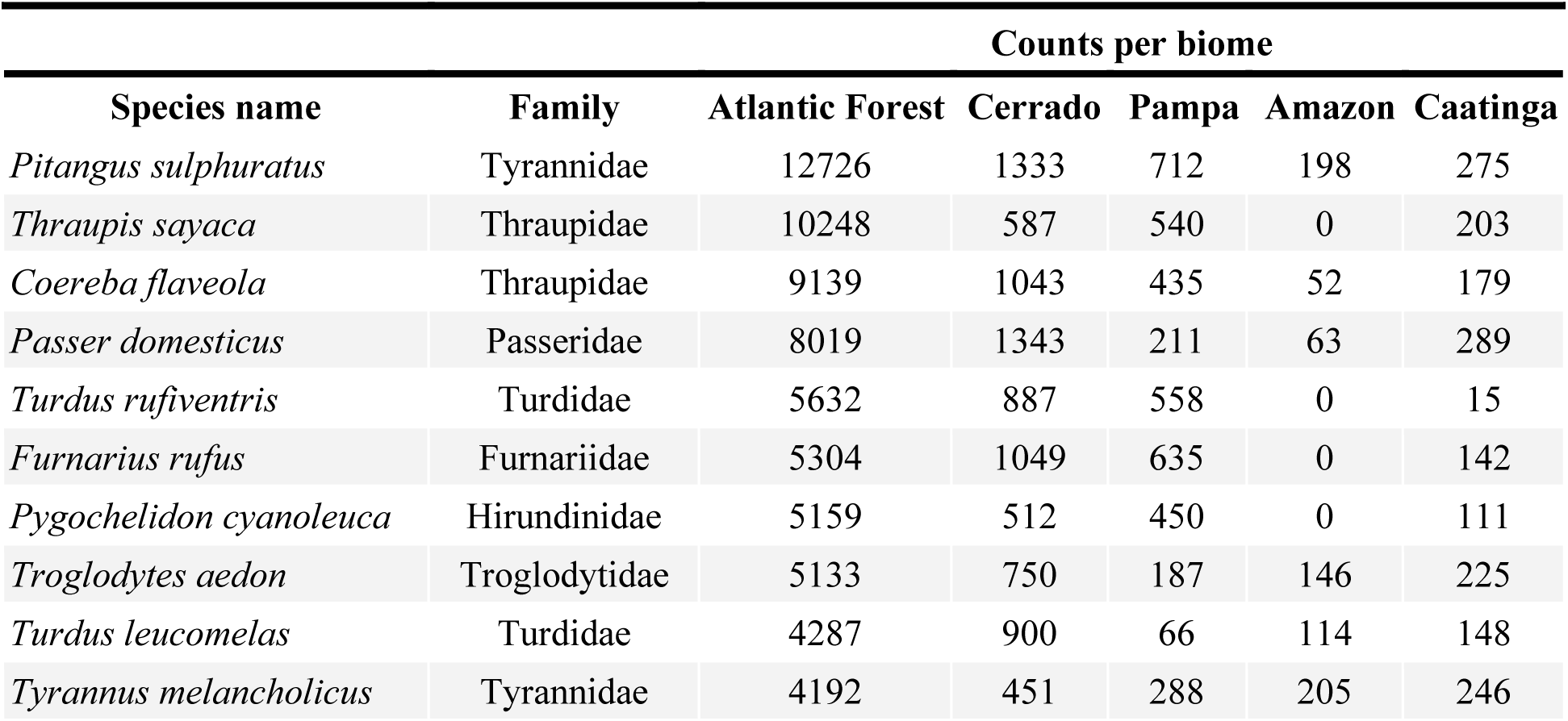
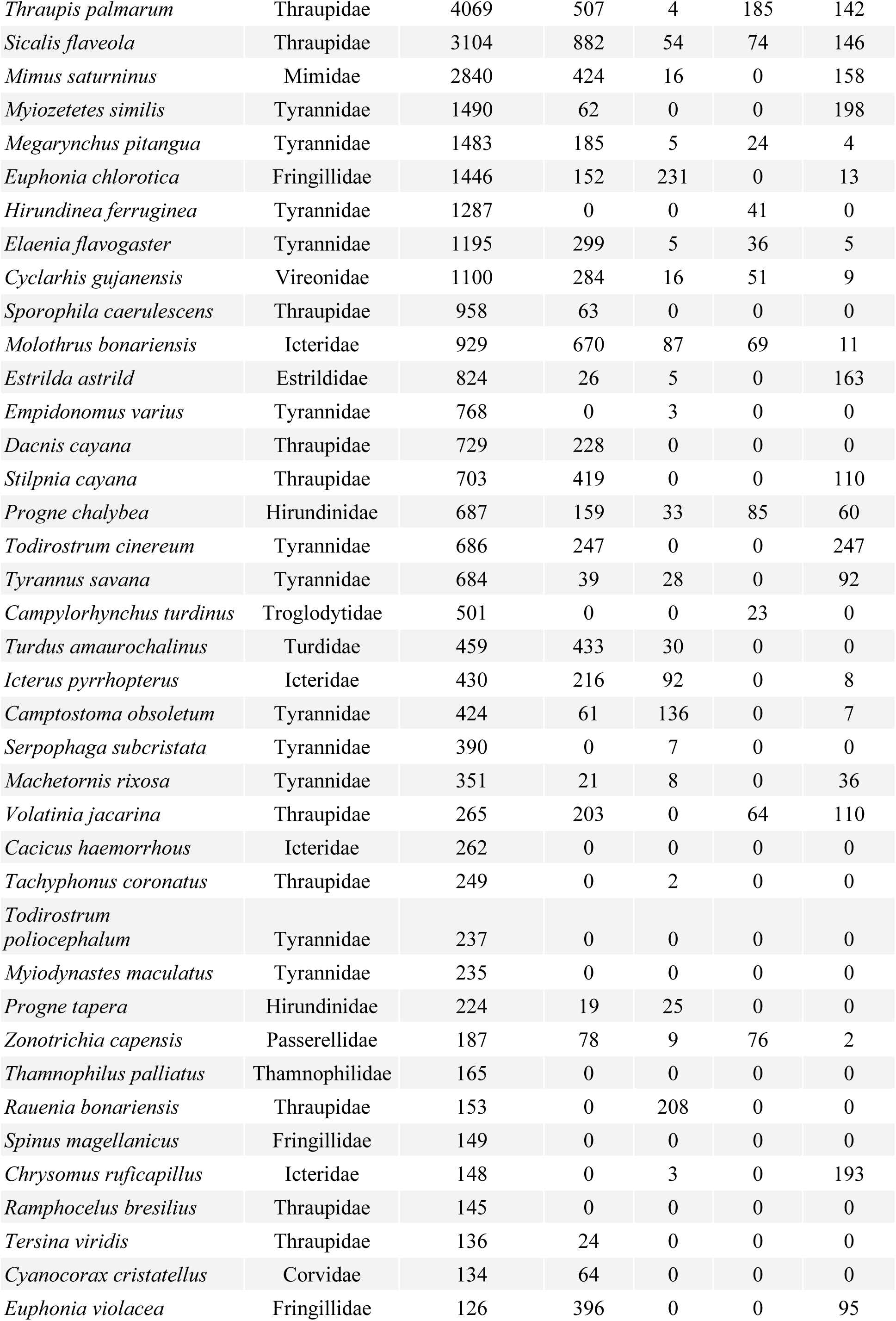
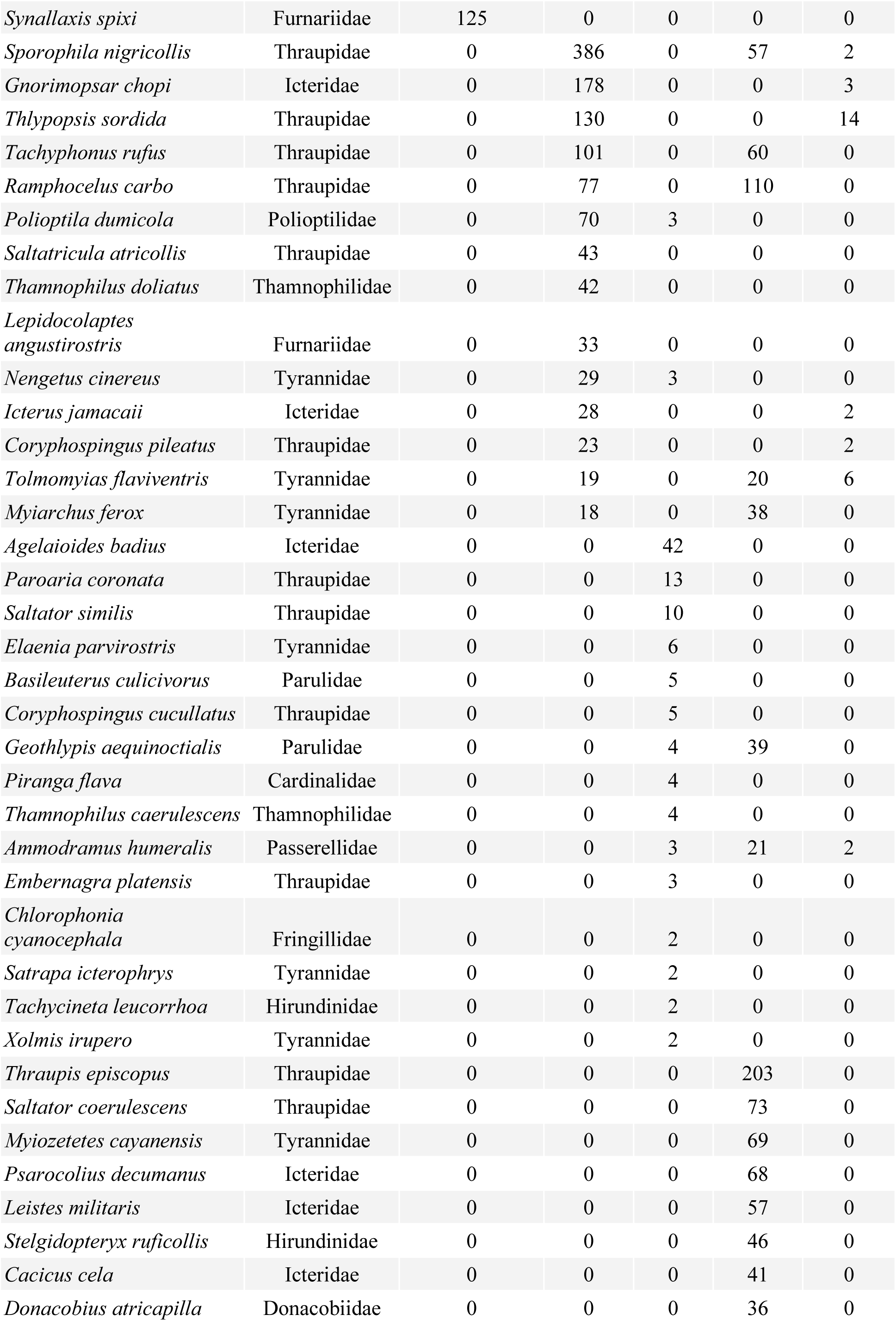
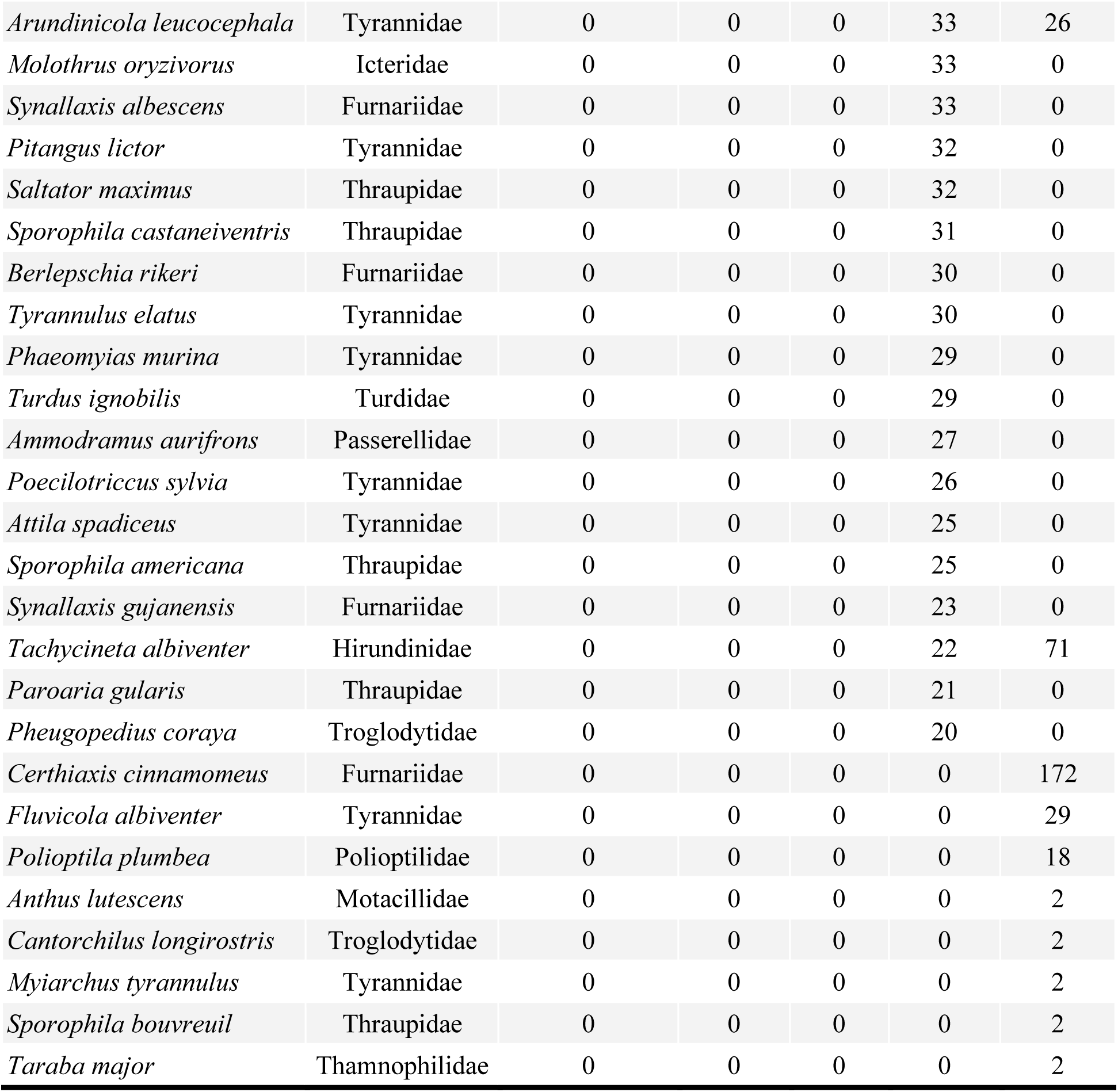
The fifth most frequently recorded species (based on the number of times they appear in checklists) in the *very high urban assemblages* per biome in the year 2021. The "very high urban assemblages" are those with a proportion of urban cells equal or greater than 75%.

## 8. References

Afrifa, Joseph K., Kweku A. Monney, and Justus P. Deikumah. "Effects of urban land-use types on avifauna assemblage in a rapidly developing urban settlement in Ghana." Urban Ecosystems 26.1 (2023): 67–79.

Aronson, M. F., Nilon, C. H., Lepczyk, C. A., Parker, T. S., Warren, P. S., Cilliers, S. S.,… & Zipperer, W. (2016). Hierarchical filters determine community assembly of urban species pools. Ecology, 97(11), 2952–2963.

Baston, D. (2022). exactextractr: Fast Extraction from Raster Datasets using Polygons, R package version 0.7.2, 2021. Retrieved from https://cran.r-project.org/web/packages/exactextractr/.

Bello, C., Galetti, M., Pizo, M. A., Magnago, L. F. S., Rocha, M. F., Lima, R. A.,… & Jordano, P. (2015). Defaunation affects carbon storage in tropical forests. Science Advances, 1(11), e1501105.

Bonier, F., Martin, P. R., & Wingfield, J. C. (2007). Urban birds have broader environmental tolerance. Biology Letters, 3(6), 670–673.

Bostwick, K. (2016). Feathers and plumages. In R. J. Lovette & J. W. Fitzpatrick (Eds.), Handbook of Bird Biology (3rd ed., pp. 101– 148). John Wiley, West Sussex, UK.

Callaghan, C. T., Major, R. E., Wilshire, J. H., Martin, J. M., Kingsford, R. T., & Cornwell, W. K. (2019). Generalists are the most urban-tolerant of birds: a phylogenetically controlled analysis of ecological and life history traits using a novel continuous measure of bird responses to urbanization. Oikos, 128(6), 845–858.

Candolin, U., & Heuschele, J. (2008). Is sexual selection beneficial during adaptation to environmental change?. Trends in Ecology & Evolution, 23(8), 446–452.

Carballo, L., Delhey, K., Valcu, M., & Kempenaers, B. (2020). Body size and climate as predictors of plumage colouration and sexual dichromatism in parrots. Journal of Evolutionary Biology, 33(11), 1543–1557.

Clements, J. F., P. C. Rasmussen, T. S. Schulenberg, M. J. Iliff, T. A. Fredericks, J. A. Gerbracht, D. Lepage, A. Spencer, S. M. Billerman, B. L. Sullivan, and C. L. Wood. (2023). The eBird/Clements checklist of Birds of the World: v2023, Cornell University, New York, USA.

Cooney, C. R., He, Y., Varley, Z. K., Nouri, L. O., Moody, C. J., Jardine, M. D.,… & Thomas, G. H. (2022). Latitudinal gradients in avian colourfulness. Nature Ecology & Evolution, 6(5), 622–629.

Cuthill, I. C., Allen, W. L., Arbuckle, K., Caspers, B., Chaplin, G., Hauber, M. E.,… & Caro, T. (2017). The biology of color. Science, 357(6350), eaan0221.

Dale, J., Dey, C. J., Delhey, K., Kempenaers, B., & Valcu, M. (2015). The effects of life history and sexual selection on male and female plumage colouration. Nature, 527(7578), 367–370.

Delhey, K., & Peters, A. (2017). Conservation implications of anthropogenic impacts on visual communication and camouflage. Conservation Biology, 31(1), 30–39.

Dunn, R. R., Gavin, M. C., Sanchez, M. C., & Solomon, J. N. (2006). The pigeon paradox: dependence of global conservation on urban nature. Conservation Biology, 1814–1816.

Evans, K. L., Hatchwell, B. J., Parnell, M., & Gaston, K. J. (2010). A conceptual framework for the colonisation of urban areas: the blackbird Turdus merula as a case study. Biological Reviews, 85(3), 643–667.

Ferreiro-Arias, I., Santini, L., Sagar, H. S. C., Richard-Hansen, C., Guilbert, E., Forget, P. M.,… & Benítez-López, A. (2024). Drivers and spatial patterns of avian defaunation in tropical forests. Diversity and Distributions, e13855, 1–15.

Giraudeau, M., Toomey, M. B., Hutton, P., & McGraw, K. J. (2018). Expression of and choice for condition-dependent carotenoid-based color in an urbanizing context. Behavioral Ecology, 29(6), 1307–1315.

Gao, J., & O’Neill, B. C. (2020). Mapping global urban land for the 21st century with data-driven simulations and Shared Socioeconomic Pathways. Nature Communications, 11(1), 2302.

Gong, P., Li, X., Wang, J., Bai, Y., Chen, B., Hu, T.,… & Zhou, Y. (2020). Annual maps of global artificial impervious area (GAIA) between 1985 and 2018. Remote Sensing of Environment, 236, 111510.

Götmark, F., & Post, P. (1996). Prey selection by sparrowhawks, *Accipiter nisus*: relative predation risk for breeding passerine birds in relation to their size, ecology and behaviour. Philosophical Transactions of the Royal Society of London. Series B: Biological Sciences, 351(1347), 1559–1577.

Grimm, N. B., Faeth, S. H., Golubiewski, N. E., Redman, C. L., Wu, J., Bai, X., & Briggs, J. M. (2008). Global change and the ecology of cities. Science, 319(5864), 756–760.

Gutiérrez-Tapia, P., Azócar, M. I., & Castro, S. A. (2018). A citizen-based platform reveals the distribution of functional groups inside a large city from the Southern Hemisphere: e-Bird and the urban birds of Santiago (Central Chile). Revista Chilena de Historia Natural, 91(3), 1–16.

Haddou, Y., Mancy, R., Matthiopoulos, J., Spatharis, S., & Dominoni, D. M. (2022). Widespread extinction debts and colonization credits in United States breeding bird communities. Nature Ecology & Evolution, 6(3), 324–331.

Hahs, A. K., & Evans, K. L. (2015). Expanding fundamental ecological knowledge by studying urban ecosystems. Functional Ecology, 29(7), 863–867.

Hahs, A. K., Fournier, B., Aronson, M. F., Nilon, C. H., Herrera-Montes, A., Salisbury, A. B.,… & Moretti, M. (2023). Urbanisation generates multiple trait syndromes for terrestrial animal taxa worldwide. Nature Communications, 14(1), 4751.

Halfwerk, W., Blaas, M., Kramer, L., Hijner, N., Trillo, P. A., Bernal, X. E.,… & Ellers, J. (2019). Adaptive changes in sexual signalling in response to urbanization. Nature Ecology & Evolution, 3(3), 374–380.

Hartig, F. (2022). DHARMa: Residual Diagnostics for Hierarchical (Multi-Level/Mixed) Regression Models. R package version 0.4.6. Retrieved from https://CRAN.R-project.org/package=DHARMa.

Hijmans, R. (2022). terra: Spatial data analysis. R package version 1.6-7. Retrieved from https://cran.r-project.org/web/packages/terra/index.html

Hilal, M., Joly, D., Roy, D., & Vuidel, G. (2018). Visual structure of landscapes seen from built environment. Urban Forestry & Urban Greening, 32, 71–80.

Hoang, N. T., & Kanemoto, K. (2021). Mapping the deforestation footprint of nations reveals growing threat to tropical forests. Nature Ecology & Evolution, 5(6), 845–853.

Iglesias-Carrasco, M., Duchêne, D. A., Head, M. L., Møller, A. P., & Cain, K. (2019). Sex in the city: Sexual selection and urban colonization in passerines. Biology Letters, 15(9), 20190257.

Janas, K., Gudowska, A., & Drobniak, S. M. (2024). Avian colouration in a polluted world: a meta-analysis. Biological Reviews. Early View.

Jetz, W., Thomas, G. H., Joy, J. B., Hartmann, K., & Mooers, A. O. (2012). The global diversity of birds in space and time. Nature, 491(7424), 444–448.

Johnston, A., Hochachka, W. M., Strimas-Mackey, M. E., Ruiz Gutierrez, V., Robinson, O. J., Miller, E. T.,… & Fink, D. (2021). Analytical guidelines to increase the value of community science data: An example using eBird data to estimate species distributions. Diversity and Distributions, 27(7), 1265–1277.

Kark, S., Iwaniuk, A., Schalimtzek, A., & Banker, E. (2007). Living in the city: can anyone become an ‘urban exploiter’? Journal of Biogeography, 34(4), 638–651.

Leveau, L. M. (2019). Urbanization induces bird color homogenization. Landscape and Urban Planning, 192, 103645.

Leveau, L. (2021). United colours of the city: A review about urbanisation impact on animal colours. Austral Ecology, 46(4), 670–679.

Lim, J. Y., Svenning, J. C., Göldel, B., Faurby, S., & Kissling, W. D. (2020). Frugivore-fruit size relationships between palms and mammals reveal past and future defaunation impacts. Nature Communications, 11(1), 4904.

McKinney, M. L. (2006). Urbanization as a major cause of biotic homogenization. Biological Conservation, 127(3), 247–260.

Morellato, L. P. C., & Haddad, C. F. (2000). Introduction: The Brazilian Atlantic Forest 1. Biotropica, 32(4b), 786–792.

Naimi, B., Hamm, N. A., Groen, T. A., Skidmore, A. K., & Toxopeus, A. G. (2014). Where is positional uncertainty a problem for species distribution modelling? Ecography, 37(2), 191–203.

Neate-Clegg, M. H., Tonelli, B. A., Youngflesh, C., Wu, J. X., Montgomery, G. A., Şekercioğlu, Ç. H., & Tingley, M. W. (2023). Traits shaping urban tolerance in birds differ around the world. Current Biology, 33(9), 1677–1688.

Partecke, J., Van’t Hof, T., & Gwinner, E. (2004). Differences in the timing of reproduction between urban and forest European blackbirds (Turdus merula): result of phenotypic flexibility or genetic differences?. Proceedings of the Royal Society of London. Series B: Biological Sciences, 271(1552), 1995–2001.

Patankar, S., Jambhekar, R., Suryawanshi, K. R., & Nagendra, H. (2021). Which traits influence bird survival in the city? A review. Land, 10(92), 1–22.

Pebesma, E.; Bivand, R. (2023). Spatial Data Science: With Applications in R (1st ed.). 314 pages. Chapman and Hall/CRC, Boca Raton. 10.1201/9780429459016

Pocock, M. J., Tweddle, J. C., Savage, J., Robinson, L. D., & Roy, H. E. (2017). The diversity and evolution of ecological and environmental citizen science. PLoS ONE, 12(4), e0172579.

R Core Team (2023). R: A Language and Environment for Statistical Computing. R Foundation for Statistical Computing, Vienna, Austria. https://www.R-project.org/

Renoult, J. P., Kelber, A., & Schaefer, H. M. (2017). Colour spaces in ecology and evolutionary biology. Biological Reviews, 92(1), 292–315.

Senior, R. A., Oliveira, B. F., Dale, J., & Scheffers, B. R. (2022). Wildlife trade targets colorful birds and threatens the aesthetic value of nature. Current Biology, 32(19), 4299–4305.

Seto, K. C., Güneralp, B., & Hutyra, L. R. (2012). Global forecasts of urban expansion to 2030 and direct impacts on biodiversity and carbon pools. Proceedings of the National Academy of Sciences, 109(40), 16083–16088.

Sol, D., González-Lagos, C., Moreira, D., Maspons, J., & Lapiedra, O. (2014). Urbanisation tolerance and the loss of avian diversity. Ecology Letters, 17(8), 942–950.

Sol, D., Bartomeus, I., González-Lagos, C., & Pavoine, S. (2017). Urbanisation and the loss of phylogenetic diversity in birds. Ecology Letters, 20(6), 721–729.

Souza Jr, C. M., Z. Shimbo, J., Rosa, M. R., Parente, L. L., A. Alencar, A., Rudorff, B. F.,… & Azevedo, T. (2020). Reconstructing three decades of land use and land cover changes in brazilian biomes with landsat archive and earth engine. Remote Sensing, 12(17), 2735.

Stoddard, M. C., & Prum, R. O. (2008). Evolution of avian plumage color in a tetrahedral color space: a phylogenetic analysis of new world buntings. The American Naturalist, 171(6), 755–776.

Strimas-Mackey, M. et al. Best Practices for Using eBird Data. Version 1.0. https://cornelllabofornithology.github.io/ebird-bestpractices/. (Cornell Lab of Ornithology, 2020) https://doi.org/10.52

Sullivan, B. L., Wood, C. L., Iliff, M. J., Bonney, R. E., Fink, D., & Kelling, S. (2009). eBird: A citizen-based bird observation network in the biological sciences. Biological Conservation, 142(10), 2282–2292.

Tribot, A. S., Deter, J., & Mouquet, N. (2018). Integrating the aesthetic value of landscapes and biological diversity. Proceedings of the Royal Society B: Biological Sciences, 285(1886), 20180971.

Tobias, J. A., Sheard, C., Pigot, A. L., Devenish, A. J., Yang, J., Sayol, F.,… & Schleuning, M. (2022). AVONET: morphological, ecological and geographical data for all birds. Ecology Letters, 25(3), 581–597.

Turak, N., Monnier-Corbel, A., Gouret, M., & Frantz, A. (2022). Urbanization shapes the relation between density and melanin-based colouration in bird communities. Oikos, 2022(12), e09313.

Zhao, M., Cheng, C., Zhou, Y., Li, X., Shen, S., & Song, C. (2022). A global dataset of annual urban extents (1992–2020) from harmonized nighttime lights. Earth System Science Data, 14(2), 517–534.

Yeh, P. J. (2004). Rapid evolution of a sexually selected trait following population establishment in a novel habitat. Evolution, 58(1), 166–174.

Yu, J., Duan, H., Zhang, B., Zhang, L., & He, J. (2024). Urbanization alters the geographic patterns of passerine plumage color in China. Landscape and Urban Planning, 248, 105101.

Wickham, H., Averick, M., Bryan, J., Chang, W., McGowan, L. D. A., François, R.,… & Yutani, H. (2019). Welcome to the Tidyverse. Journal of Open Source Software, 4(43), 1686.

